# A new pathway in central metabolism mediates nutrient control of development and antibiotic production by *Streptomyces*

**DOI:** 10.1101/2024.07.14.603434

**Authors:** Chao Li, Mia Urem, Ioli Kotsogianni, Josephine Lau, Somayah S. Elsayed, Nathaniel I. Martin, Iain W. McNae, Patrick Voskamp, Christoph Mayer, Sébastien Rigali, Navraj Pannu, Jan Pieter Abrahams, Lennart Schada von Borzyskowski, Gilles P. van Wezel

## Abstract

The amino sugar *N-*acetylglucosamine (GlcNAc) plays a central role in primary metabolism and is a key signaling molecule for the onset of morphological and chemical differentiation of *Streptomyces*. The global nutrient-sensory regulator DasR acts as the gatekeeper of development in streptomycetes, and its activity is modulated by aminosugar phosphates. Here, we report the discovery of a novel pathway in aminosugar metabolism that governs GlcNAc sensing. GlcNAc-6P is converted into a toxic metabolite via two new enzyme functions, namely dehydration of *N*-acetylglucosamine-6-phosphate by NagS to form 6P-Chromogen I, a reaction that has not yet been described in the textbooks, and its subsequent deacetylation by NagA producing a cytotoxic structural analogue of ribose. The latter reveals an unexpected promiscuous activity for GlcNAc-6P deacetylase NagA. The crystal structures of NagS apoenzyme and NagS in complex with its substrate GlcNAc-6P or its inhibitor 6-phosphogluconate were resolved at 2.3 Å, 2.6 Å, and 1.7 Å resolution, respectively. Detailed *in vivo* and *in vitro* studies resolved the key residues of the NagS catalytic site. Thus, we have uncovered a novel pathway in aminosugar metabolism that sheds new light on nutrient-mediated control of development and antibiotic production in *Streptomyces*.

## INTRODUCTION

Amino sugars are important nutrients for bacteria. *N*-acetylglucosamine (GlcNAc) is the monomer of chitin, the second most abundant polysaccharide on earth after cellulose, and widely distributed in both soil and aqueous environments^1–4^. GlcNAc is also part of the bacterial peptidoglycan (PG), which consists of chains of alternating GlcNAc and *N-*acetylmuramic acid (MurNAc) residues cross-linked via peptide bridges^5^. GlcNAc is a preferred carbon and nitrogen source for streptomycetes, and their metabolism is biased for the used of this sugar ^6^. Streptomycetes are Gram-positive filamentous bacteria with a complex multicellular life cycle^7^. They are known as nature’s medicine makers, producing two thirds of all clinical antibiotics, as well as many other natural products with bioactivity, such as anticancer, anthelmintic, antifungal and immunosuppressant^8–10^. During the onset of development, an autolytic process results in the degradation of the cell wall in the substrate mycelium. This process provides nutrients for building the aerial mycelium, which eventually develop into spores^11,12^. GlcNAc plays a key role in nutrient sensing and response, regulating the initiation of development and antibiotic production. Under rich nutritional conditions (*feast*), GlcNAc activates growth, thereby repressing development and antibiotic production. Conversely, under poor growth conditions (*famine*), GlcNAc activates both processes^13^.

Metabolic control of development is mediated via the GntR-family regulator DasR^13,14^. DasR is a global nutrient sensory regulator that controls the uptake and metabolism of GlcNAc, and is involved in the regulation of all pathway-specific regulators for antibiotic and siderophore biosynthesis in *Streptomyces coelicolor* ^15–17^. GlcNAc-6P and GlcN-6P allosterically induce the release of DasR from its recognition sites (**Fig. S1**)^13,17–19^. GlcNAc is imported by the phosphoenolpyruvate phosphotransferase system (PTS), internalized as GlcNAc-6P^6,20^. GlcNAc-6P deacetylase NagA converts GlcNAc-6P into GlcN-6P, which is then deaminized by GlcN-6P deaminase NagB to produce the glycolytic intermediate fructose-6P (Fru-6P). Thus, GlcN-6P stands at the crossroads of multiple key metabolic pathways, namely glycolysis via conversion to Fru-6P by NagB and peptidoglycan synthesis following its conversion to UDP-GlcNAc^21–23^ (**Fig. S1**). It is important to note – and at the same time surprising - that GlcN does not act as signaling molecule for development, even though it only requires a single deacetylation to convert GlcNAc-6P into GlcN-6P. Hence, GlcN and GlcNAc play distinct role in the *Streptomyces* life cycle, which so far has not been explained.

Accumulation of high concentrations of phosphorylated aminosugars is toxic to bacteria, as shown for *Bacillus subtilis* and *Escherichia coli* ^2^ and for *Streptomyces*^24^. *S. coelicolor nagB* mutants are sensitive to both GlcN and GlcNAc, and a lethal screen resulted in spontaneous Δ*nagB* suppressor mutants that could grow in the presence of either GlcNAc or GlcN^24^. However, GlcN and GlcNAc toxicity appear to act via independent pathways, as several suppressors of GlcN toxicity still fail to grow on GlcNAc and *vice versa*. Recently, we published that deletion of the gene for the ROK-family regulator RokL6 relieves GlcN toxicity, which is entirely due to the enhanced expression of the MFS transporter SCO1448^25^. SCO1448 likely exports a toxic compound, but cannot relieve the toxicity of GlcNAc. This again shows major differences in the perception of GlcN and GlcNAc by streptomycetes.

In this work, we show that mutation or deletion of the gene for SCO4393, which contains a sugar isomerase (SIS) domain, blocks GlcNAc sensing and relieves its toxicity to *nagB* mutants. The enzyme was renamed NagS, for *<u>N</u>*-<u>a</u>cetyl<u>g</u>lucosamine sensitivity. NagS remarkably acts as an *N*-acetylglucosamine-6-phosphate dehydratase, an activity that has never been described in any organism. NagA deacetylates the product of NagS to form a ribose-like compound that is key to GlcNAc toxicity. We also identified a likely salvage pathway, whereby the pentose phosphate pathway intermediate 6-phosphogluconate acts as a metabolic inhibitor of NagS, which means that enhanced flux of PPP reduces GlcNAc-6P toxicity in wild-type cells. Crystal structures were resolved for NagS apoenzyme and for NagS in complex with its substrate GlcNAc-6P or its metabolic inhibitor 6-phosphogluconate. Our data provide insights into a unique metabolic pathway within streptomycetes that explains nutrient-mediated control of morphological and chemical differentiation.

## RESULTS

### NagS is required for GlcNAc toxicity and aminosugar sensing

Mutants of *Streptomyces coelicolor* that lack *nagB* fail to grow on minimal media (MM) supplemented with GlcNAc. In order to obtain mutants that are affected in GlcNAc sensing and nutrient control of development and antibiotic production, we analyzed mutants obtained in a suppressor screen, selecting for spontaneous mutants of *S. coelicolor nagB* mutants that survive specifically on GlcNAc. Sequencing of suppressor mutant SM11 revealed a single nucleotide polymorphism (SNP) at nucleotide position 535 of SCO4393 (G to A substitution), leading to a non-silent change from aspartate to asparagine (D179N) in the predicted gene product. SCO4393 contains a so-called SIS domain^26^, which spans the entire protein (see below).

To verify the role of NagS in GlcNAc metabolism and nutrient sensing, we created a *nagB*-*nagS* double mutant. For this, the coding region of *nagS* from positions +15 to + 768 was removed from the *S. coelicolor nagB* mutant to generate an in-frame deletion mutant (see **Extended Methods**). While the parental strain *S. coelicolor* M145 grew normally on MM with 1% (w/v) mannitol and 10 mM GlcNAc, the *nagB* mutant was unable to grow under these conditions. Conversely, both the suppressor mutant SMA11 and the Δ*nagB*Δ*nagS* mutant grew well in the presence of GlcNAc. To ascertain that the deletion of *nagS* was the sole cause of the observed phenotypes described, we genetically complemented the Δ*nagB*Δ*nagS* mutant with a native copy of *nagS*, whereby the - 451/+908 region of *nagS* was amplified from *S. coelicolor* chromosome and cloned into pSET152, generating strain Δ*nagB*Δ*nagS*^C^. Expression of *nagS* restored sensitivity of the mutants to GlcNAc (**Fig. 1a**). These data strongly suggest that the deletion of *nagS* was the sole cause of the restored growth on GlcNAc.

**Figure 1.**
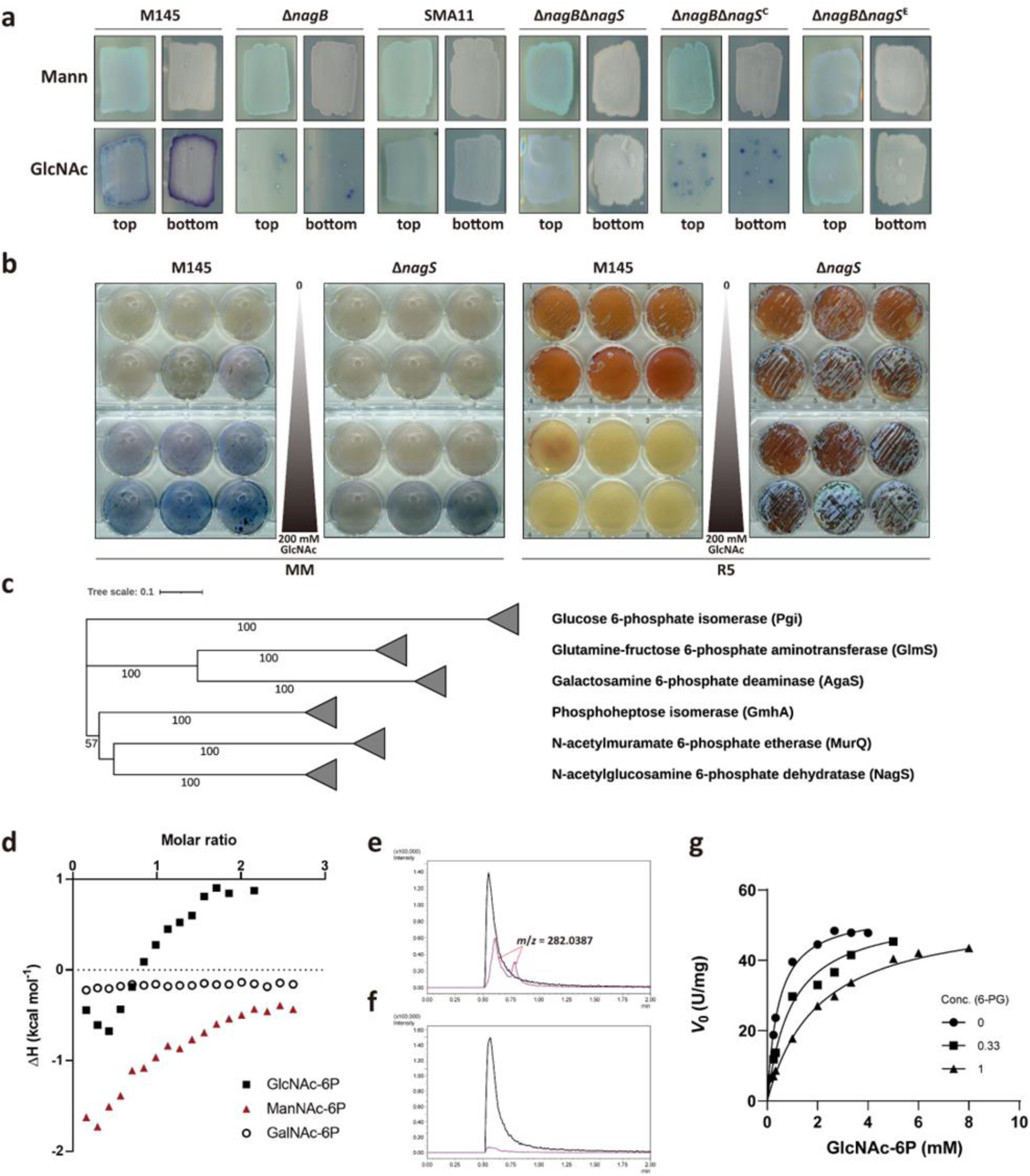
NagS is a novel GlcNAc-6P dehydratase. (**a**) Sensitivity of *S. coelicolor* mutants to GlcNAc. Spores (5 × 10^5^ CFU) of *S. coelicolor* M145 and its mutant derivatives Δ*nagB*, SMA11, Δ*nagB*Δ*nagS*, Δ*nagB*Δ*nagS*^C^ (Δ*nagB*Δ*nagS* expressing *nagS*) and Δ*nagB*Δ*nagS*^E^ (Δ*nagB*Δ*nagS* with empty plasmid pSET152) were streaked on MM agar plates with 1% mannitol (Mann) and 1% mannitol plus 10 mM GlcNAc (GlcNAc). Strains were cultured for 72 h at 30°C. Note that *nagB* mutants can grow only when SCO4393 (*nagS)* has been mutated (suppressor SMA11) or deleted. (**b**) NagS and its role in GlcNAc sensing. Spores of M145 and Δ*nagS* were plated on MM and R5 with 0, 0.001, 0.01, 0.1, 1, 5, 10, 20, 50, 100, 150, 200 mM GlcNAc. Note that *nagS* mutants hardly respond to GlcNAc. (**c**) Phylogenetic tree of several different types of sugar isomerase (SIS) domain enzymes in bacteria, including NagS, Glucose-6P isomerase (Pgi), Phosphoheptose isomerase (GmhA), *N*-acetylmuramic acid-6-phosphate etherase (MurQ), Glutamine-fructose-6-phosphate aminotransferase (GlmS), and putative D-galactosamine-6-phosphate deaminase (Agas). The phylogenetic tree was made by MEGA11 (Neighbour-joining method) and built based on the alignment of the amino acid sequences. (**d**) ITC analysis of NagS with *N*-acetyl-6-phosphate amino sugar metabolites, namely GlcNAc-6P, ManNAc-6P, and GalNAc-6P. Both GlcNAc-6P and ManNAc-6P are bound well by NagS, while the enzyme did not bind to GalNAc-6P. (**e-f**) confirmation of NagS products by LC-MS. Extracted ion chromatograms for GlcNAc-6-P (black trace) and its dehydrated product (pink trace) in the enzymatic reaction mixture of GlcNAc-6-P with either active NagS (**e**) or heat-inactivated NagS (**f**). Peaks relating to the product (*m*/*z* = 282.0387) are indicated by red arrows. (**g**) Kinetics of NagS in the presence of 6-PG (0, 0.33, 1 mM). The *V*_0_ data used in **d** were plotted against the substrate concentration, and each assay was performed in triplicate and expressed as a mean ± standard error.

We then wondered if NagS would play a role in GlcNAc-dependent control of development and specialized metabolism. For this, *S. coelicolor* was cultured on minimal media (MM) and nutrient-rich R5 agar plates with increasing concentrations of GlcNAc. Expectedly, GlcNAc enhanced development and antibiotic production of the wild-type *S. coelicolor* on MM and inhibited these processes on R5. In contrast, the *nagS* mutant had largely lost GlcNAc sensing, showing normal sporulation on R5 agar even at GlcNAc concentrations up to 200 mM, and the mutant failed to show enhanced antibiotic production when MM was supplemented with GlcNAc (**Fig. 1b**). Furthermore, we previusly showed that besides development and antibiotic production, GlcNAc also blocks siderophore production by *S. coelicolor* M145 on R5 agar plates. We therefore wondered if the biosynthesis of siderophores would still be subjected to catabolite repression by GlcNAc. Interestingly, *nagS* null mutants had lost the ability to suppress siderophore production on R5 with added GlcNAc (**Fig. S2**). Taken together, these observations suggest that NagS plays a crucial role in amino sugar sensing in streptomycetes. This suggests that NagS plays an essential role in nutrient dependent control of differentiation.

Next, we constitutively expressed either *nagA* or *nagS* in the *S. coelicolor nagB* null mutant using the *ermE* promoter, to see if we could still obtain suppressor mutants. Expectedly, constitutive expression of either *nagA* or *nagS* in Δ*nagB* reduced the appearance of suppressor mutants when the strains were grown on GlcNAc, because in that case two active copies of either *nagA* or *nagS* are present in the cells (**Fig. S3a**). However, expression of either NagA in the *nagB*-*nagS* double mutant or NagS in the *nagAB* double mutant, did not prevent normal growth on MM with 10 mM GlcNAc (**Fig. S3b**). These data show that both NagA and NagS are required to cause GlcNAc toxicity, strongly suggesting a link between the two metabolic activities.

### The phylogenetically conserved NagS uses GlcNAc-6P and ManNAc-6P as substrates

NagS contains a conserved SIS domain^26^, predicted to span almost the entire length of the protein. SIS domains are typically found in phosphosugar-binding proteins, including *N*-acetylmuramic acid-6-phosphate etherase (MurQ) and glutamine-fructose-6-phosphate aminotransferase (GlmS)^27–29^. Phylogenomic analysis revealed that NagS is conserved in all streptomycetes, with high amino acid sequence similarity between the predicted gene products (**Fig. S4a**). Gene synteny analysis showed that the genomic region around *nagS* is also conserved, whereby *nagS* is invariably located adjacent to and divergently expressed from the iron master regulatory gene *dmdR1* (SCO4394). Furthermore, linkage between *nagS*-*dmdR1* exists in all *Streptomycetaceae* except for the genus *Yinghuangia* (**Fig. S4b**). NagS is closely related to MurQ, but forms a well-defined and separate clade in the phylogenetic tree (**Fig. 1c**), suggesting that the protein may have a function distinct from the other enzymes.

In order to identify the substrate(s) of NagS, we first expressed NagS-His_6_ in *E. coli* and purified the protein to homogeneity using Ni-affinity chromatography. Size-exclusion chromatography revealed it to be a dimer. Isothermal titration calorimetry (ITC) was performed with glucose-6-phosphate (Glc-6P), Fru-6P, GlcN-6P, GlcNAc, GlcNAc-1P and GlcNAc-6P. Of these, only GlcNAc-6P showed significant binding to NagS (**Fig. S4c**), suggesting that both the *N*-acetyl moiety and the C6 phosphate group are important for NagS binding. We then tested binding of GlcNAc-6P and its epimers *N*-acetylmannosamine-6-phosphate (ManNAc-6P) and *N*-acetylgalactosamine-6-phosphate (GalNAc-6P) as ligands. This revealed that both GlcNAc-6P and ManNAc-6P were bound by NagS, while GalNAc-6P was not (**Fig. 1d**). Notably, a change in absorbance at 230 nm was detected when either 2 mM GlcNAc-6P or ManNAc-6P were incubated with purified NagS at 30 °C (**Fig. S4d,e**), suggesting that NagS converted both compounds *in vitro* into hitherto unknown products.

### NagS is an *N*-acetylglucosamine 6-phosphate dehydratase

The product formed from the conversion of GlcNAc-6P by NagS was identified based on NMR and LC-MS analysis. Following *in vitro* reactions using GlcNAc-6P as the substrate, new peaks were detected in the ^1^H NMR spectrum specifically for the reaction with active NagS, which were not seen following incubation with heat-inactivated enzyme (**Fig. S5a**). The chemical shifts of the reaction product pointed at the formation of phosphorylated Chromogen I, the 2,3-dehydro derivative of GlcNAc (**Table S4**, **Fig. S6**)^30^. 6P-Chromogen I (2-acetamido-2,3-dideoxy-6-phosphate-D-erythro-hex-2-enofuranose; designated **1**) exists as a mixture of its α and β anomers. This was further confirmed by LC-MS analysis, which showed the appearance of two peaks in the reaction mixture containing the active NagS (**Fig. 1e,f**). The HRESIMS spectrum of the peaks (**Fig. S5b**) established a molecular formula of C_8_H_14_NO_8_P (*m/z* 282.0387), which is consistent with the structure identified by NMR for **1** (**Table S5**). NagS also acts on the other substrate, ManNAc-6P, resulting in its dehydration, although the NMR signals corresponding to **1** were relatively lower in intensity as compared to the reaction with GlcNAc-6P (**Fig. S5c**). Kinetics demonstrated a markedly higher catalytic efficiency of NagS for GlcNAc-6P, with *k*and *K* values of 24.67 s^−1^ and 0.45 mM, respectively, which is approximately 40 times greater than that for ManNAc-6P, measured at 0.90 s^−1^ and 0.68 mM, respectively (**Table 1** and **Fig. S5e,f**).

**Table 1.** Kinetic parameters for NagS.

| NagS* | GlcNAc-6P | ManNAc-6P |
| --- | --- | --- |
| $K_m$ (mM) | $0.45 \pm 0.03$ | $0.68 \pm 0.05$ |
| $k_{cat}$ ($s^{-1}$) | $24.67 \pm 0.38$ | $0.90 \pm 0.02$ |
| $k_{cat}/K_m$ ($M^{-1} \cdot s^{-1}$) | $5.48 \times 10^4$ | $1.32 \times 10^3$ |

\* Each assay was carried out in triplicate and is expressed as mean ± standard error.

Taken together, our data show that NagS dehydrates both GlcNAc-6P and ManNAc-6P to produce **1** (**Fig. S5d**). NagS and its homologues constitute a completely novel family of GlcNAc-6P dehydratases within the SIS superfamily.

### 6-phosphogluconate is a competitive inhibitor of NagS

Next, we sought to identify potential other interaction partners of NagS to better understand its role in *S. coelicolor* carbon metabolism. Since 6-phosphogluconate (6-PG) structurally resembles linear GlcNAc-6P (**Fig. S7a**), we assessed the interaction between 6-PG and NagS using thermal denaturation assays (see **Extended Methods**). Interestingly, the addition of 1 mM 6-PG shifted the *T*_m_ of NagS from 48.84°C ± 0.10°C and 59.53°C ± 0.31°C, to 51.74°C ± 0.58°C and 57.91°C ± 0.58°C; while the addition of 5 mM 6-PG shifted the *T*_m_ from two separate peaks to a single merged peak with *T*_m_ at 53.95°C ± 0.20°C when compared against the NagS control (**Fig. S7a**). 6-PG was not metabolized when incubated with NagS (**Fig. S7b**), but instead it effectively inhibited the conversion of GlcNAc-6P (**Fig. S7c**). Further kinetic analysis revealed that while the *V*_max_ remained constant, the *K*_m_ values for GlcNAc-6P increased with rising concentrations of 6-PG, observed at 0.33 mM and 1 mM (**Fig. 1g** and **Fig. S7d**). The inhibition constant (*K*_i_) for 6-PG was determined to be 0.28 ± 0.03 mM, supporting its role as a competitive inhibitor. Additionally, the sensitivity of Δ*nagB* to GlcNAc was mitigated by supplementation with exogenous D-gluconate (**Fig. S7e**).

### The crystal structure of NagS

The atomic structures of apo-NagS, and its complexes with GlcNAc-6P and 6-PG, were determined by X-ray crystallography to resolutions of 2.3, 2.6 and 1.7 Å, respectively (**Table S3**). The structures revealed that the biologically active dimer of NagS interacts along a crystallographic twofold axis with C2 symmetry (**Fig. 2a**). Two identical substrate binding cavities are located at the dimeric interface (**Fig. S8a**). This dimer interface, which sandwiched the substrate, is stabilized by evolutionarily conserved salt bridges and hydrogen bonds. Structurally, each monomer adopts an α/β configuration with a three-layered αβα sandwich architecture, featuring a standard parallel β-sheet surrounded by nine α-helices. The central β-sheet consists of five strands (β1-β5), contributing to the stability and functionality of the enzyme (**Fig. 2b**).

**Figure 2.**
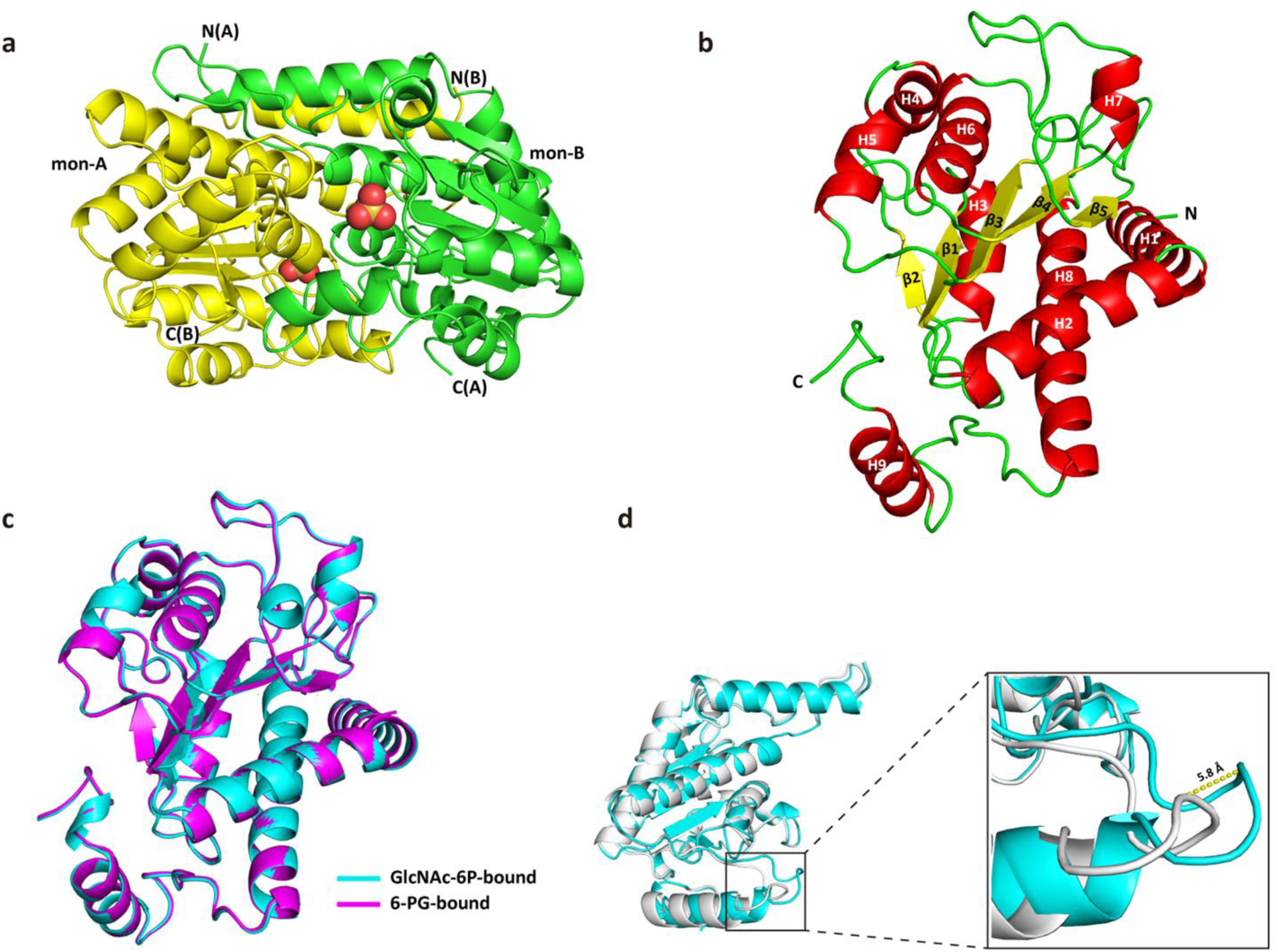
Crystal structure of NagS. (**a**) Dimeric structure of NagS determined by X-ray crystallography. The individual monomers (mon-A and mon-B) of the dimer are shown in green and yellow, with the N- and C-terminus indicated. Sulphates are shown in spheres. (**b**) Secondary structure of monomeric NagS. α-helices are shown in red, while β-sheets are shown in yellow and loops are shown in green. Monomer of NagS contains nine α-helices, H1-H9, and five β-sheets, β1-β5, as indicated. (**c**) Alignment of the secondary structure of GlcNAc-6P bound NagS monomer (cyan) with that of monomer NagS in complex with 6-PG (magenta), with an RMS deviation of 0.17 for all atoms. (**d**) Alignment of the secondary structure of NagS apo monomer (white) with that of monomer NagS in complex with GlcNAc-6P (cyan), with an RMS deviation of 0.22 (200 to 200 atoms). The significant change of the loop (residues 226-232) is shown in the red box and zoomed in. The loop of NagS in complex with GlcNAc-6P shows a 5.8 Å shift as indicated by the yellow dotted line.

The protein structures of GlcNAc-6P-, and 6-PG-bound NagS were virtually indistinguishable from each other with an RMS deviation of only 0.17 for all atoms (**Fig. 2c**). However, the apo-form deviated from the substrate-bound structures. In the substrate-bound conformations, the loops connecting H8 and H9 (residues 226-232) had closed in on the substrate (**Fig. 2d**, **Fig. S8b**), resulting in a maximum main chain shift of 5.8 Å at the Cα of Val229. These loops are located at opposite sides of the dimer, and no other structural shifts were observed, indicating that substrate binding is unlikely to be collaborative. The function of the movable loop is therefore most likely to ensure a compact complex between NagS and GlcNAc-6P, and to trigger catalysis by bringing some of its catalytic side chain residues in contact with the substrate.

### Analysis of the GlcNAc-6P binding site

Clear electron density at the NagS catalytic site demonstrates that residues from both monomers collaborate to form a tight substrate-binding pocket (**Fig. 3a,b**). Adjacent to this site, a well-ordered water molecule, likely involved in the initial ring-opening step of catalysis, is observed (**Fig. 3c**). Additionally, several other well-ordered water molecules are integral to the active site structure. The phosphate groups of GlcNAc-6P and 6-PG engage Ser54, Ser119, and Ser121 from one monomer, resembling the phosphate-binding mechanism seen in MurQ, where three serine residues play a similar crucial role in phosphate binding^31,32^. The GlcNAc moiety is stabilized through interactions with Ser54, Ser91, and Glu94, as well as the main chain amide of His53 from the same monomer. Furthermore, it interacts with the side chains of ArgB64 and AsnB228, and the main chain amides of AlaB65 and GlyB227 from the adjacent monomer. Similar interactions are found in the 6-PG-bound state (**Fig. 3d,e**).

**Figure 3.**
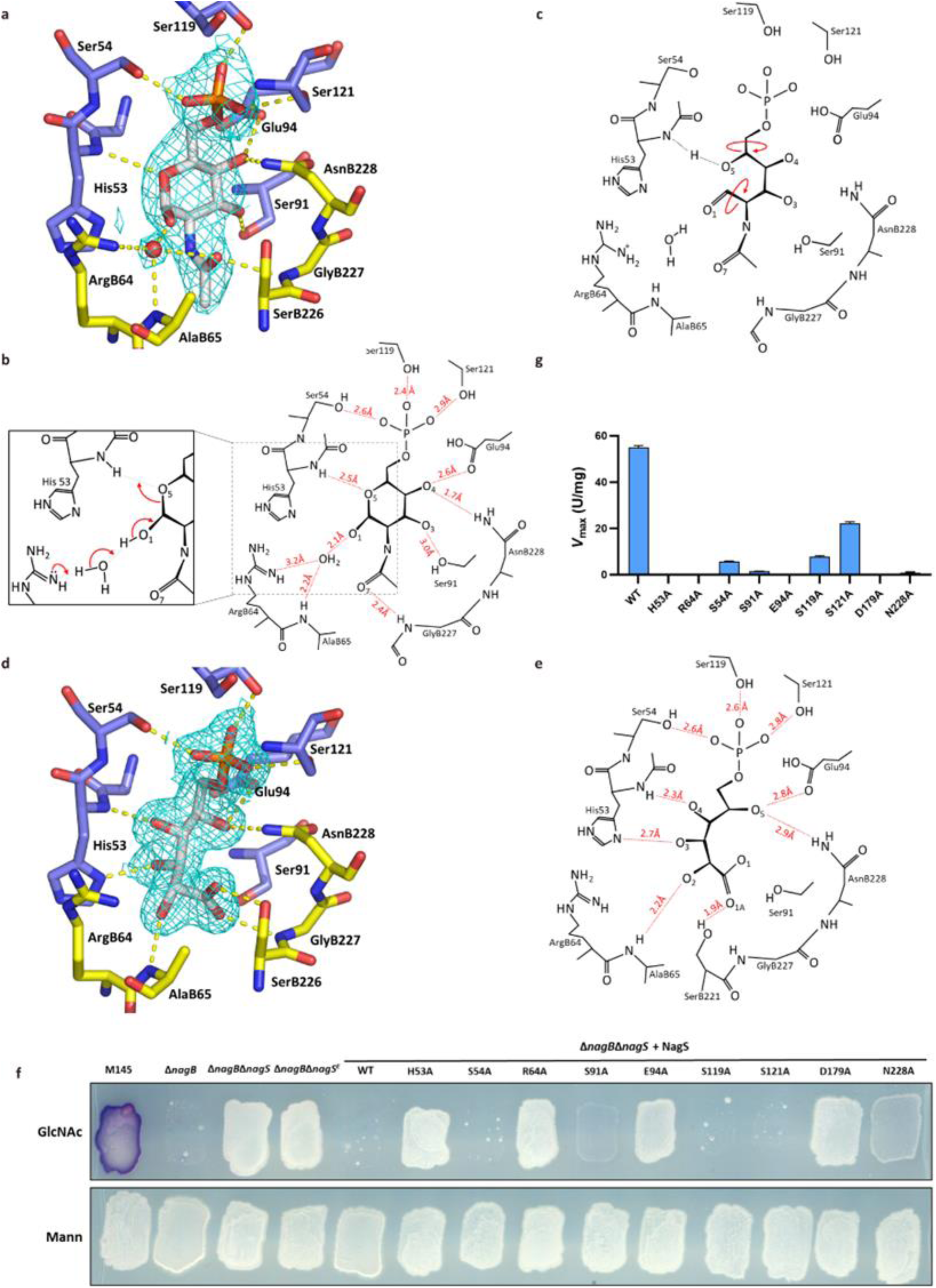
Analysis of the binding site of NagS and activity of NagS mutants. (**a**) NagS active site with bound GlcNAc-6P (grey carbons), protein residues are coloured with pale blue and yellow carbons to indicate the 2 monomers forming the active site. 2|Fo|-|Fc| electron density contoured at 1.2 σ is shown as cyan mesh. Hydrogen bonds are indicated by dashed yellow lines. (**b**) GlcNAc-6P binding site of NagS, with hydrogen bonding distances, or distances between hydrogens and hydrogen bond acceptors indicated. The other molecule of the dimer contributes aminoacids ArgB64, AlaB65, GlyB227 and AsnB228. The inset indicates the likely electron rearrangements required for ring opening, the first step of catalysis. (**c**) Putative transition state after ring opening, prior to rotations about the C5-C6 and C1-C2 bonds of GlcNAC-6P (indicated in red), that presumably precede subsequent ring closing. These rotations are associated with the rearrangement of hydrogen bonds. This likely requires conformational changes that in the crystal are inhibited by crystal contacts, explaining why the crystals are not enzymatically active. (**d**) NagS active site with bound 6-phosphogluconate (grey carbons), protein residues are coloured with pale blue and yellow carbons to indicate the 2 monomers forming the active site. 2|Fo|-|Fc| electron density contoured at 1.2 σ is shown as cyan mesh. (**e**) Hydrogen bonding distances observed in the 6-phosphogluconic-inhibited state of NagS. The inhibited state suggests how GlcNAc-6P rearranges upon ring-opening, and likely reflects the transition state prior to ring closing, which probably involves Ser91, Glu94, and AsnB228. (**f**) NagS enzyme activity *in vivo*. GlcNAc sensivity of Δ*nagB*Δ*nagS* harbouring clones expressing NagS mutants H53A, S54A, R64A, S91A, E94A, S119A, S121A, D179A or N228A were grown on MM agar supplemented with 1% mannitol (Mann) and 1% mannitol plus 10 mM GlcNAc (GlcNAc). Single colonies are most likely spontaneous suppressors. (**g**) *In vitro* activity (*V*_max_) for wild-type NagS (WT) and NagS variants, with the substrate of GlcNAc-6P.

To investigate the roles of specific residues in binding and catalysis, alanine mutants of all residues that interact with GlcNAc-6P through their side chains together with Asp179, which was determined to be essential for NagS activity from the suppressor mutant, were generated and expressed in the *S. coelicolor* double mutant Δ*nagB*Δ*nagS*. The phenotypes of these mutants were analyzed on MM agar plates, with and without GlcNAc. Expression of wild-type *nagS* or its mutant variants, S54A, S119A, or S121A, fully restored GlcNAc sensitivity in the double mutant, indicating that these residues are not crucial for the catalytic activity of NagS. Conversely, mutants S91A and N228A only partially restored GlcNAc sensitivity, underscoring their relevance in the enzyme’s catalytic function. However, constructs expressing the H53A, R64A, E94A, or D179A mutants of NagS showed no restoration of GlcNAc sensitivity, indicating that these residues are essential for NagS activity (**Fig. 3f**).

To corroborate these findings, we determined the kinetic parameters of all of the NagS variants *in vitro*, which aligned closely with our *in vivo* observations (**Table 2** and **Fig. S9**). Variants S54A, S119A, and S121A displayed relatively small changes in *K*_m_ and *k*_cat_ values, compared to the other mutants, confirming these residues are not crucial for catalysis. In contrast, variants S91A and N228A exhibited significantly reduced activity, with very low *V*_max_ values (**Fig. 3g**). Specifically, S91A demonstrated a 40-fold decrease in *k*_cat_/*K*_m_, while N228A showed a 208-fold decrease in *k*_cat_/*K*_m_ and a 3.9-fold reduction in *K*_m_, indicating that they are important for catalysis. Furthermore, the variants H53A, R64A, E94A and D179A were completely inactive (**Table 2** and **Fig. 3g**), confirming that these residues are critical for catalysis.

**Table 2.** Kinetic parameters for NagS variants.

| NagS variants* | $K_m$ (mM) | $k_{cat}$ ($s^{-1}$) | $k_{cat}/K_m$ ($M^{-1} \cdot s^{-1}$) |
| --- | --- | --- | --- |
| H53A<sup>#</sup> | - | - | - |
| S54A | $0.85 \pm 0.09$ | $2.70 \pm 0.09$ | $3.30 \times 10^3$ |
| R64A<sup>#</sup> | - | - | - |
| S91A | $0.56 \pm 0.04$ | $0.78 \pm 0.01$ | $1.38 \times 10^3$ |
| E94A<sup>#</sup> | - | - | - |
| S119A | $2.06 \pm 0.27$ | $3.83 \pm 0.23$ | $1.86 \times 10^3$ |
| S121A | $1.19 \pm 0.09$ | $10.49 \pm 0.28$ | $8.84 \times 10^3$ |
| D179A | - | - | - |
| N228A | $1.77 \pm 0.28$ | $0.47 \pm 0.03$ | 263.30 |

\* The substrate tested was GlcNAc-6P; Each assay was carried out in triplicate and is expressed as mean ± standard error.
<sup>#</sup> "-" indicates that no reaction was observed.

### A novel function for NagA as 6P-chromogen I deacetylase

The observation that the deletion of *nagA* relieves the toxicity of GlcNAc to *nagB* null mutants created a mystery in terms of the enzyme’s role in GlcNAc metabolism. NagS is the first step towards a toxicity pathway, and in the absence of NagA the NagS substrate GlcNAc-6P will accumulate in large amounts when cells grow on GlcNAc, which should therefore be more, rather than less, toxic to the cells. However, *nagA nagB* double mutants grow very well. We therefore wondered if NagA might have a second yet unknown enzymatic function. The most obvious way for NagA to prevent toxicity would be if besides GlcNAc-6P it also deacetylates 6P-chromogen I, the reaction product of NagS. To test this idea, *S. coelicolor* NagA was expressed in *E. coli* and purified to homogeneity, and its activity with different substrates was tested. NagA deacetylated GlcNAc-6P, ManNAc-6P and GalNAc-6P *in vitro*, with a clear preference for GlcNAc-6P (*k* /*K* of 1.01 × 10^5^ M^−1^·s^−1^) as compared to 6.06 × 10^3^ M^−1^·s^−1^ for ManNAc-6P and 1.03 × 10^4^ M^−1^·s^−1^ for GalNAc-6P (**Table 3** and **Fig. S10a-c**). These kinetic parameters are comparable to those for NagA isozymes reported in *E. coli* and *Mycobacterium tuberculosis*^33,34^, which shows that the purified NagA was fully active *in vitro*. Next, we tested if NagA could act on **1**. GlcNAc-6P was first incubated with NagS at 30°C; next, either active NagA or heat-inactivated NagA was added and the reaction mixture analysed by LC-MS. No difference in peak intensity was seen when heat-inactivated NagA was added, showing that **1** was stable under the given reaction conditions. Excitingly, **1** disappeared after adding active NagA, and new products with an exact mass of 240.0280 (**Table S5**) were detected in the reaction mixture (**Fig. 4a,b**). This shows that NagA is a promiscuous enzyme, whereby besides its textbook function it can also deacetylate **1**, the product of NagS. The HRESIMS spectrum of the peaks established a molecular formula of C_6_H_12_NO_7_P (*m/z* 240.0280), which is the deacetylated product of **1** (**Fig. S10d**).

**Figure 4.**
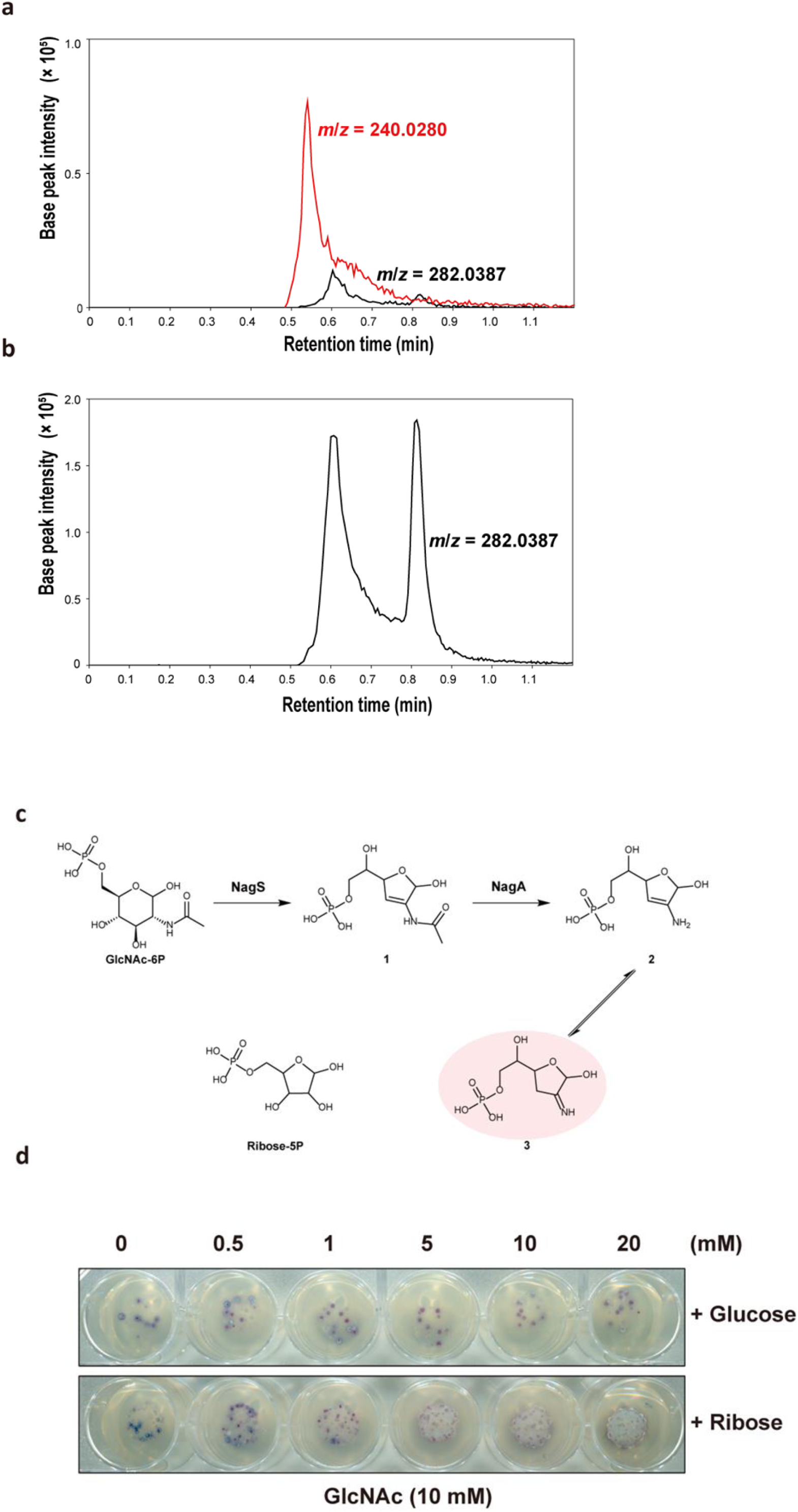
New function of *S. coelicolor* NagA and an updated aminosugar metabolic pathway. Extracted ion chromatograms for compound **1** (*m*/*z* 282.0387, shown in black lines) and compound **3**/**4** (*m*/*z* 240.0280, shown in red lines) in the enzymatic reaction mixture of GlcNAc-6P with NagS and NagA (**a**) or deactivated NagA (**b**). (**c**) Updated metabolic pathway of aminosugar in *Streptomyces*. Based on the metabolic pathway in **Fig. S1**, we propose a new metabolic way in GlcNAc metabolism. Apart from the canonical reaction whereby GlcNAc-6P is metabolised by NagA and NagB to fructose-6P, GlcNAc-6P can also be dehydrated by NagS and subsequently deacetylated by NagA to form compound **3** (shown in light red), whose is a likely toxic compound whose chemical structure is similar to ribose-5P. (**d**) Ribose by-passes GlcNAc toxicity. Spores (5 × 10^5^ CFU) of the *S. coelicolor* M145 *nagB* mutant were spotted on MM supplemented with 1% mannitol plus 10 mM GlcNAc and different concentrations (0 to 20 mM) of either glucose or D-ribose, followed by incubation for 72 h at 30°C. Note that 1mM ribose or more alleviates GlcNAc toxicity and allows the cells to grow, while even at 20 mM glucose the colonies still are sensitive to GlcNAc. Single colonies are most likely suppressors.

**Table 3.** Kinetic parameters for *S. coelicolor* NagA.

| NagA* | GlcNAc-6P | ManNAc-6P | GalNAc-6P |
| --- | --- | --- | --- |
| $K_m$ (mM) | $1.59 \pm 0.21$ | $2.40 \pm 0.44$ | $2.84 \pm 0.41$ |
| $k_{cat}$ ($s^{-1}$) | $159.61 \pm 9.02$ | $14.56 \pm 1.10$ | $29.17 \pm 2.31$ |
| $k_{cat}/K_m$ ($M^{-1} \cdot s^{-1}$) | $1.01 \times 10^5$ | $6.06 \times 10^3$ | $1.03 \times 10^4$ |

\* Each assay was carried out in triplicate and is expressed as mean ± standard error.

Thus, our work reveals a novel metabolic route in central aminosugar metabolism (**Fig. 4c**). Herein, GlcNAc-6P is dehydrated by NagS to **1**. Subsequently, **1** is deacetylated by NagA to form enamine **2** (2-amino-2,3-dideoxy-6-phosphate-D-erythro-hex-2-enofuranose), which is unstable and spontaneously converts to the corresponding imine **3** (2-imino-2,3-dideoxy-6-phosphate-D-erythro-hexofuranose).

### Supplementation of ribose relieves GlcNAc toxicity to *nagB* mutants

We then wondered what the basis might be for the toxicity of compound **3**. We noticed that imine **3** shows significant structural similarities to ribose-5-phosphate, an intermediate in pentose metabolism and the sugar moiety of nucleic acids. This suggests that **3** may interfere with the synthesis of nucleotides when accumulating at high concentrations. If this is the case, high concentrations of ribose may relieve the toxicity. Therefore, either ribose or glucose was added to cultures of the *S. coelicolor nagB* mutant grown in the presence of GlcNAc, to determine if these sugars could alleviate the toxicity of GlcNAc. Importantly, *nagB* mutants were significantly less sensitive to GlcNAc when grown in the presence of higher concentrations of ribose, while glucose did not alter the sensitivity (**Fig. 4d**). This provided an important clue as to the toxicity of **3**, which may act by interfering with ribose metabolism, and hence with the synthesis of nucleotides.

## DISCUSSION

*N*-acetylglucosamine (GlcNAc) is a preferred carbon source for streptomycetes and stands at the crossroads of aminosugar metabolism, glycolysis and cell wall synthesis. The molecule also plays a key role in nutrient sensing and the ultimate developmental decision to initiate sporulation and antibiotic production^13^. GlcNAc inhibits development under rich growth conditions (*feast*), while it activates development and antibiotic production under poor growth conditions (*famine*)^13^. Our work uncovered a previously unknown route in central metabolism, namely a toxicity pathway that is mediated by GlcNAc sensing in streptomycetes, governed by NagS and NagA. Deletion of the gene for NagS annihilates GlcNAc signaling, with *nagS* mutants showing normal development on both MM and R5 agar, regardless of how much GlcNAc is added to the plates. Additionally, deletion of *nagS* anihilated the effect of GlcNAc on specialized metabolites, exemplified by both the pigmented antibiotics Act and Red, as well as siderophores. NagS and NagA form a pathway in central metabolism that cannot be found in current textbooks (see e.g. http://www.kegg.jp/). Discovery of one new enzyme function in central metabolism is rare, let alone two that form a novel pathway.

GlcNAc-6P dehydratase NagS converts GlcNAc-6P to 6P-Chromogen I (**1**), which is subsequently deacetylated via a previously unknown promiscuous enzymatic activity of NagA to enamine **2**, which spontaneously converts to the corresponding imine **3** (**Fig. 4c**). The reaction catalyzed by NagS most likely initiates with a dehydration step that involves opening the glucose 6-ring, followed by its closure into a 2’,3’-dideoxy-2’,3’-unsaturated ribose 5-ring. This transformation most likely begins with the ring-opening of the glucose moiety in GlcNAc-6P, and we propose this process mirrors the ring-opening mechanism of glucose-6-phosphate by phosphoglucose isomerase^35^. ArgB64 in NagS likely assumes a role similar to Lys518 in phosphoglucose isomerase through a coordinating a water molecule (**Fig. 3c**). The acidic Asp179 then forms a salt bridge with ArgB64. Its mutation to Asparagine or Alanine would prevent the guanidinium group of ArgB64 from properly orienting to attack O1 of the GlcNAc-6P substrate. This ring-opening likely precipitates a series of rotations around single bonds within the carbohydrate, aligning it into a configuration similar to that of the linear inhibitor 6-PG. However, 6-PG is distally stabilized through an interaction between its O1A and SerB221’s side chain, an interaction unlikely to occur with the linearized GlcNAc-6P due to steric hindrance caused by its longer carbohydrate chain. We hypothesize that subsequent bond formation between O1 and C4, coupled with dehydrolysis removing O3 and forming a double bond between C2 and C3, is facilitated by Glu94, with the support of Ser91 and AsnB228. Although our data do not conclusively detail these latter steps, the successful capture of GlcNAc-6P bound to the active site in pre-formed crystals indicates that substrate reorientation—necessitating rotations around single bonds that precede ring-closing and dehydrolysis—likely requires conformational rearrangements of the protein, which were impeded by crystal contacts.

An important question is what the basis is for aminosugar toxicity, which has been reported in several bacteria ^2,24^. The fact that compound **3** structurally resembles ribose-5P provided a clue as to the nature of the toxicity. Indeed, addition of ribose relieved the toxicity of GlcNAc to *nagB* mutants, which suggests that the toxicity pathway governed by NagS and NagA might interfere with nucleic acid synthesis. Lysis of the vegetative or substrate mycelia is a key event driving colony growth and development, through a process of programmed cell death (PCD). Streptomycetes that produce larger concentrations of cytotoxic compounds with antiproliferative activity such as prodiginines^12^ or anthracyclines^36^ are strongly impaired in development. Prodiginines play a role in the onset of PCD in *S*. *coelicolor* as DNA and membrane damaging agent, killing biomass in the centre of the colony to provide nutrients and allowing the colony to expand^12^. Activation of prodiginine production is mediated by GlcNAc by direct expression control by DasR of the *red* biosynthetic gene cluster^37^. Moreover, it was also shown that the onset of prodiginine production coincided in time and space with an area of the *S*. *coelicolor* culture with high density of dying filaments^38^. Our work thus suggests that the area of mycelium undergoing cell death and triggering prodiginine production may well result – at least in part – from the accumulation of the toxic compound associated with GlcNAc metabolism and generated by NagS and NagA.

Loss of siderophore-mediated iron import is another key factor in GlcNAc-mediated developmental arrest. On nutrient-rich R5 agar, GlcNAc inhibits siderophore production and therefore iron import by metabolically inhibiting DasR; this relieves its repression of *dmdR1*, which encodes the iron homeostasis repressor in *Streptomyces*. The GlcNAc-mediated activation of *dmdR1* expression thus results in decreased desferrioxamine E (DFOE) biosynthesis, which is crucial for development of *S*. *coelicolor*^39,40^. Addition of iron to R5 agar with GlcNAc restores antibiotic production and sporulation in *S. coelicolor*^39^ which suggests that the imported Fe^2+^ also contributes to cell death prior to development, by generating reactive oxygen species (ROS) by the Fenton reaction and/or as cofactor of essential developmental genes/proteins. Gene synteny analysis shows that *nagS* lies immediately adjacent to and divergently expressed from *dmdR1* in all streptomycetes, providing strong phylogenetic support for linkage between NagS and iron homeostasis. Furthermore, repression of siderophore production by GlcNAc is lost in *nagS* null mutants. These data directly link NagS, GlcNAc metabolism, DasR and iron accumulation with PCD and the onset of morphological and chemical differentiation.

Combining all data, we propose a new model for the control of development and antibiotic production in streptomycetes, wherein accumulation of the toxic compound produced by NagS and NagA from GlcNAc-6P plays a key role (**Fig. 5**). The concentration of GlcN(Ac)-6P is thereby pivotal. As vegetative growth progresses, the substrate mycelia are degraded to feed the new biomass. This will lead to the accumulation of building blocks, such as amino acids (from proteins), nucleotides (from DNA and RNA) and GlcNAc (from the cell wall). The accumulation of GlcNAc would first participate in dismantling of the substrate mycelium via a PCD-like process, by directly activating the NagS/NagA toxicity pathways and prodiginine production. GlcNAc concentrations will rapidly increase in the part that undergoes extensive lysis, and DasR will therefore be metabolically inhibited by GlcNAc-6P and GlcN-6P, thus allowing DmdR1 to become active. As a result, iron import is reduced, thereby limiting cell death associated with Fe^2+^-mediated ROS production. How then does NagS become inactivated? We propose that like for DasR, this happens at the posttranslational level, via a *salvage pathway* that leads to metabolic inhibition of NagS and thus stops the production of the toxic compound (**Fig. 5**). A major clue came from the discovery that the pentose phosphate pathway (PPP) intermediate 6-phosphogluconate is the inhibitor of NagS (**Fig. 1g, Fig. 3d,e**). The compound can be derived in a few steps from GlcN-6P, by its conversion to fructose-6P by NagB, then to glucose-6P by phosphoglucose isomerase (Pgi), and to 6-PG via the PPP. It is important to note that this also explains why *nagB* mutants are sensitive to GlcN-6P or GlcNAc-6P; the cells cannot convert GlcN-6P into Fru-6P, thus blocking the salvage pathway, explaining the high sensitivity of *nagB* mutants to GlcNAc. GlcNAc would therefore participate in processes either activating or repressing PCD.

**Figure 5.**
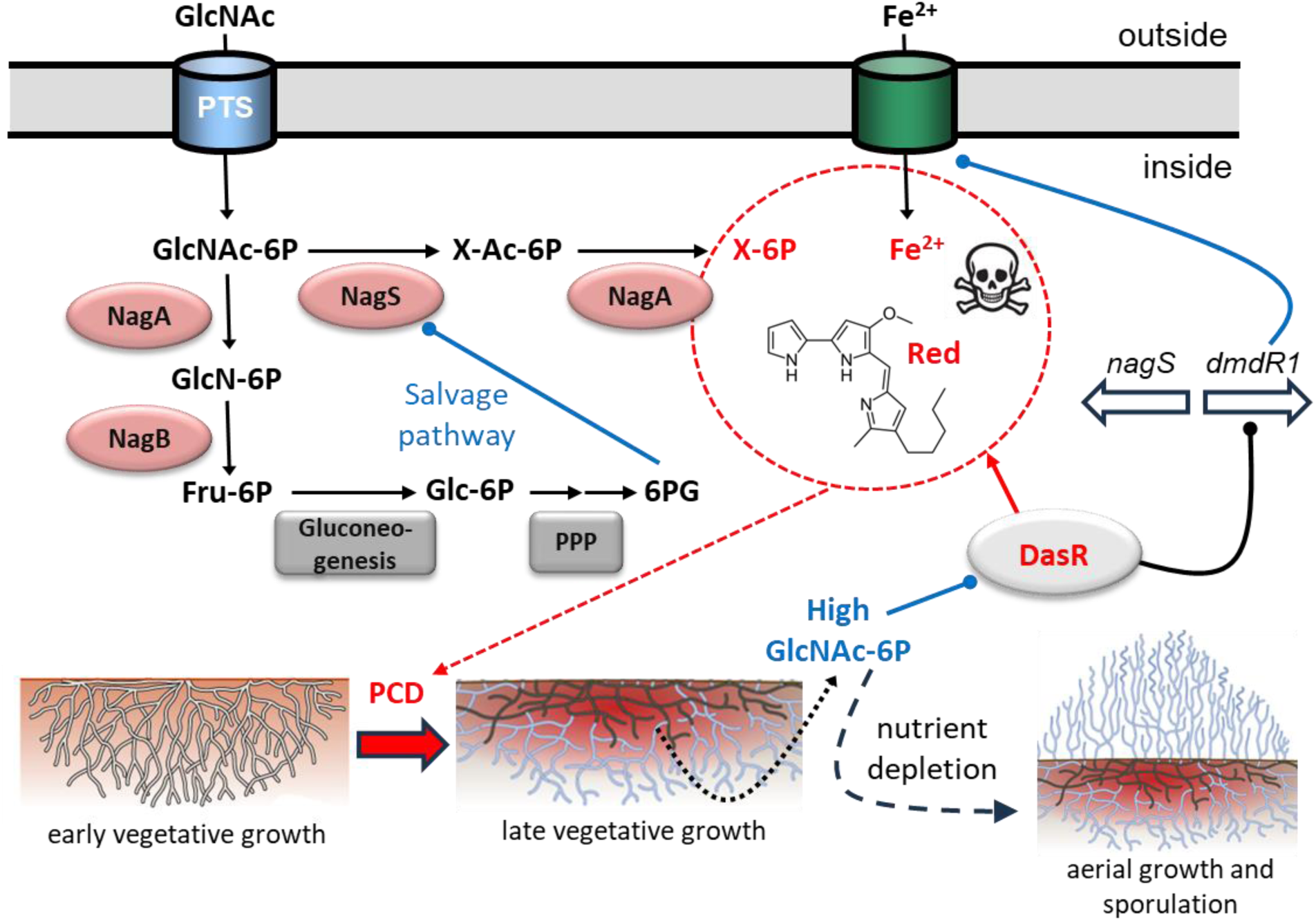
Model for the metabolic control of development by GlcNAc under nutrient-rich conditions. Red indicates toxicity mechanisms, in blue mechanisms that counteract toxicity. During vegetative growth, three toxic molecules accumulate, namely (i) iron imported by siderophores; (ii) prodiginines; and (iii) the toxic metabolite (denoted as X-6P) generated by NagS and NagA; this promotes PCD, a process that dismantles the old vegetative or substrate hyphae. During PCD, development is postponed, promoting expansion of the colony to provide sufficient biomass prior to the onset of development ^12^. Mycelial lysis leads to increased accumulation of GlcNAc, which is imported by the PTS. High GlcN(Ac)-6P concentrations inhibit DasR, and DmdR1 is expressed, which represses iron import and prodiginine production ceases. The salvage pathway leads to accumulation of 6-phosphogluconate (6PG), which inhibits NagS. In this way, all three toxicity mechanisms cease, thus ending the PCD phase. The GlcNAc-6P pool is gradually consumed, and nutrient depletion triggers development. The production of toxic compound X-6P from GlcNAc by NagS and NagA is key to this developmental block. For details see the text.

By counteracting accumulation of said cytotoxic molecules, PCD will come to an end. Nutrient depletion is a major signal for the onset of morphological and chemical differentiation^41,42^; GlcNAc-6P will be gradually consumed, and once levels are low again, aerial growth is initiated. Finally, it is important to note that in experiments on agar plates the pool of GlcNAc (some 20 ml of agar containing 10 mM GlcNAc) can never be depleted by the colonies, so that development remains suppressed, explaining the *feast* phenotype in the laboratory, where development is permanently blocked.

In conclusion, we report on the function and structure of the aminosugar dehydratase NagS, a novel enzyme associated with GlcNAc metabolism in *Streptomyces* that together with NagA produces a toxic compound. We thereby extend the current knowledge of central metabolism, and shed new light on the basis of aminosugar toxicity. Accumulation of the metabolite generated from the new aminosugar pathway formed by NagS-NagA explains how the onset of development and antibiotic production in streptomycetes are mediated by GlcNAc.

## METHODS

### Heterologous expression and purification of NagS

For *in vitro* enzyme experiments and structure elucidation via X-ray crystallography, N-terminally His_6_-tagged NagS and its variants with site-directed mutations (H53A, S54A, R64A, S91A, E94A, S119A, S121A, D179A and N228A) were expressed in *E. coli* Rosetta™(DE3)pLysS (Novagen). Briefly, *nagS* coding sequence was PCR amplified from *S. coelicolor* genomic DNA, while its site-directed mutated sequences were amplified from the vectors for mutated NagS complementation (pCOM-4393-H53A to pCOM-4393-N228A) (see **Extended Methods**). The amplified fragments were cloned into pET-15b to obtain the expressing plasmids, which were transformed into *E. coli* Rosetta™(DE3)pLysS competent cells. Expression host cells were grown at 37°C with shaking at 200 rpm in LB media supplemented with chloramphenicol (25 μg·mL^−1^) and ampicillin (100 μg·mL^−1^) until an OD_600_ of 0.6-0.8 was reached. Protein expression was induced with isopropyl β-D-1-thiogalactopyranoside to a final concentration of 0.5 mM, followed by further incubation at 37°C with shaking at 200 rpm for 4 hours. His_6_-tagged NagS and its variants were purified using a Ni-NTA column (GE Healthcare) with Isolation Buffer (500 mM NaCl, 5% glycerol, 50 mM HEPES, 10 mM β-mercaptoethanol, pH 8.0) containing 250 mM imidazole, as described^43^. Fractions containing the target proteins were pooled and concentrated before further purification by size exclusion chromatography (Superdex 200) on an AKTA Pure FPLC system (Cytiva) with Isolation Buffer. Proteins were desalted with HEPES buffer (20 mM HEPES, 300 mM NaCl, 5% glycerol, 1 mM DTT, pH 7.4) prior to use in crystallization trials and enzyme assays.

### Enzyme assays

All enzyme experiments were performed at 30°C using a Cary 60 UV-Vis spectrophotometer (Agilent). Each kinetic assay was carried out in triplicate and in quartz cuvettes (Hellma) with a path length of 10 mm. The initial velocity (*V*_0_) measurements were reproducible within 10% error. All stocks of enzymes and substrates were kept on ice during the entire experiments. The chemicals used were purchased from Sigma-Aldrich unless stated otherwise.

For NagS kinetic analysis, the rate of reaction was measured by following the increase in UV absorbance at 230 nm as a function of time. The reaction (300 μL in total) was started by the addition of the substrate (final concentration 0 - 5 mM) to a mixture containing 150 μL phosphate buffer (100 mM phosphate, 100 mM NaCl, pH 7.4) and 220 nM NagS. The extinction coefficient of the product of NagS was experimentally determined to be 2.17 × 10^3^ M^−1^·cm^−1^ (see **Extended Methods** and **Fig. S11**).

Kinetic analysis of *S. coelicolor* NagA was performed using a previously described direct continuous spectrophotometric assay^33^. Here, the rate of reaction was obtained from measurement of the decrease of the amide absorbance of GlcNAc-6P/ManNAc-6P/GalNAc-6P at 215 nm. The reaction mixture (300 μL) contained 150 μL phosphate buffer, *S. coelicolor* NagA (150 nM for GlcNAc-6P and 400 nM for ManNAc-6P and GalNAc-6P) and the substrates, and the reaction was initiated by the addition of substrate (concentration from 0.33 to 6.67 mM). The extinction coefficients were 500 M^−1^·cm^−1^ for GlcNAc-6P, 408 M^−1^·cm^−1^ for ManNAc-6P and 612 M^−1^·cm^−1^ for GalNAc-6P, all determined spectrophotometrically. For all kinetics, initial rate data were analysed and fitted with the Michaelis-Menten model using GraphPad Prism (version 8.3.0).

### Identification of the products of the enzymatic reactions

NagS product was identified by a combination of Nuclear Magnetic Resonance (NMR) and Liquid Chromatography-Mass Spectrometry (LC-MS) analysis. The reaction samples for NMR were prepared in phosphate buffer, in which 6.67 mM GlcNAc-6P/ManNAc-6P was incubated with 220 nM NagS at 30°C, and a sample with boiled NagS was used as a control. After reaching equilibrium at 10 min, the reaction mixture was freeze-dried and then dissolved in 170 μL D_2_O, and transferred to NMR tubes. The NMR spectra were acquired on a Bruker AVIII-600 spectrometer (Bruker BioSpin GmbH) at a field strength of 600 MHz. For LC-MS analysis, see **Extended Methods**.

### Structural data collection and structure determination

Purified NagS at a concentration of 15-20 mg/ml was screened for crystallization by sitting-drop vapour-diffusion using the PGA Screen (Molecular Dimensions), Clear Strategy Screens CSS-I and CSS-II (Molecular Dimensions), JCSG+ (Qiagen/Molecular Dimensions) and the PACT screen (Molecular Dimensions) as well as optimization screens at 20°C. The 75 μL reservoir of 96-well Innovaplate SD-2 plates was pipetted by a Genesis RS200 robot (Tecan) and drops were made by an Oryx6 robot (Douglas Instruments). Hexagonal NagS crystals (space group P6_5_22) were obtained from JCSG number 83 (96-well G11) which consisted of 2.0 M Ammonium sulphate, 0.1 M BIS-Tris, pH 5.5. Crystals were soaked in mother liquor with 10-20% glycerol as cryoprotectant, that included no other additives, or either 100mM of the substrate GlcNAc-6P, or 200 mM of the inhibitor 6-phosphogluconate. After loop mounting, they were flash-frozen in liquid nitrogen.

X-ray data were collected at the European Synchrotron Radiation Facility (Grenoble, France) on beamline ID-23 for the apo-enzyme: 1410 frames were collected on a Pilatus 6M detector at an X-ray wavelength of 0.9724 Angstroms, an exposure time of 0.037 seconds, transmission of 10% and an oscillation range of 0.2 degrees. NagS in complex with GlcNAc-6P data were collected on beamline ID-29 with a Pilatus 6M detector. For the native crystal, 1020 images were collected at 1.2727 Å wavelength with an exposure time of 0.02 seconds, transmission of 100% and an oscillation range of 0.05 degrees. We collected 680 frames at 0.976251 Angstroms wavelength with an exposure time of 0.02 seconds, transmission of 47.34% and an oscillation range of 0.1 degrees. XDS^44^ was used to process all the data collected. Scaling and merging were done using the CCP4 program ‘aimless.’^45^ The diffraction data of the 6-phosphogluconate inhibited crystals were collected at the Diamond synchrotron radiation facility on beamline I04. 3600 images were collected at a wavelength of 0.9 Å with an exposure of 0.05 seconds, transmission of 100% and oscillation of 0.05 degrees. Data processing was performed via Xia2^46^. The resolution of the inhibited crystal form was 1.7 Å, which was significantly higher than the apo- and substrate-bound crystals (2.3 Å and 2.6 Å, respectively). The structure of apo NagS was solved by molecular replacement using the structure of a putative phosphoheptose isomerase from *B. halodurans* C-125 (PDB code 3CVJ) as the model. Two subunits were present in the asymmetric unit, and in the initial stages of structure refinement, non-crystallographic symmetry restraints were imposed. Clear densities corresponding to well-ordered water molecules and either 6-Phosphogluconate, or GlcNAc-6P for the substrate-bound crystal forms, were observed for all three crystal forms. Models that included the inhibitor, or the substrate where appropriate, were inspected and interactively rebuilt using Coot^47,48^.

Water molecules were introduced automatically in refinement using Phenix^49^. Then, non-crystallographic symmetry restraints were removed and six consistent TLS fragments per monomer were automatically generated for the inhibited crystal form and refined by Phenix. Refinement of the apo- and substrate-bound structures benefitted from reference model torsion angle restraints provided by the inhibited structure, as judged by a significant improvement in R-free. TLS refinement using the same fragments as in the inhibited structure, improved the substrate-bound, but not the apo-structure. **Table S3** shows the data collection and refinement statistics for the data sets obtained.

### Isothermal titration calorimetry (ITC) assays

To identify the possible substrates of NagS, ITC tests were performed using a MicroCal PEAQ-ITC Automated microcalorimeter (Malvern Panalytical Ltd, Malvern, UK). A 700 µM solution of substrates (GlcNAc-6P, ManNAc-6P, or GalNAc-6P) in 20 mM Tris pH 7.4, 100 mM NaCl, 50 µM DTT, was titrated into a 50 µM enzyme preparation in the same buffer. Control titrations included the titration of buffer into enzyme and substrate into buffer. The samples were equilibrated to 25°C prior to measurement. The titrations were conducted at 25°C under constant stirring at 750 rpm. Each experiment consisted of an initial injection of 0.4 µL followed by 18 separate injections of 2.0 µL into the sample cell of 200 µL. The time delay between each injection was 180 seconds, the measurements were performed with the reference power set at 5 μcal·s^−1^ and the feedback mode set on “high”. The calorimetric data obtained were analysed using MicroCal PEAQ-ITC Analysis Software Version 1.20 (Malvern Panalytical Ltd, Malvern, UK). ITC data fitting was made based on the software’s “one set of sites” fitting model. The best fit is defined by chi-squared minimization. Thermodynamic parameters are reported as the average of three experiments with the standard deviation, unless stated otherwise.

### Bioinformatics analysis

Protein homology searches were performed using BLASTp^50^. The phylogenetic tree of different SIS containing proteins was made by MEGA11^51^. Synttax was used for gene synteny^52^. Protein alignments were analysed by Clustal Omega (www.ebi.ac.uk/Tools/msa/clustalo/) and visualised using Jalview (Version 2.11.2.7).

## Data availability

Atomic coordinates of the NagS models were deposited in the RCSB PDB under accession number 9F7O for apo-NagS, 9F7V for GlcNAc-6P-bound NagS and 9EOL for 6-PG-bound NagS.

## ACKNOWLEDGEMENTS

We thank Jackie Plumbridge and Malcolm Wilkinshaw for stimulating discussions and to Ellen de Waal and Helga van der Heul for technical assistance. We are grateful to the University of Edinburgh Centre for Translational and Chemical Biology and the Edinburgh Protein Production Facility for use of their facilities. The work was supported by ERC Advanced Grant 101055020-COMMUNITY of the European Union and by VICI grant 10379 from The Netherlands Organization for Scientific Research (NWO) to G.P.v.W. and by grant 201904910552 from the Chinese Scholarship Council (CSC) to C.L. The authors declare no conflicts of interest.

