## Supplemental Information for "A new pathway in central metabolism mediates nutrient control of development and antibiotic production by *Streptomyces*"

Belonging to the manuscript

### Extended Methods

#### Bacterial strains, culture conditions, plasmids and oligonucleotides

All strains described in this work are listed in Supplementary Table1. *Escherichia coli* was grown and transformed according to standard procedures[1], with *E. coli* JM109 serving as the host for routine cloning, and *E. coli* ET12567[2] for the isolation of non-methylated DNA for transformation into *Streptomyces*[3]. For protein heterologous expression, *E. coli* Rosetta™(DE3)pLysS was used for NagS and *E. coli* BL21(DE3) was used for NagA. *E. coli* was grown in Luria-Bertani (LB) media in the presence of selective antibiotics as required, with the following final concentrations: apramycin (50 µg/ml), kanamycin (50 µg/ml), ampicillin (100 µg/ml) and chloramphenicol (25 µg/ml).

*Streptomyces coelicolor* A(3)2 M145 was obtained from the John Innes Centre strain collection and served as the parent of all mutants. *S. coelicolor nagB* mutants, Δ*nagB*, and *nagAB* mutants, Δ*nagAB*, and *nagB* suppressor mutants SMA11, have been described previously[4,5]. All *Streptomyces* media and routine techniques, including transformation via conjugation and protoplast regeneration, are described in the *Streptomyces* manual[3]. A mixture of 1:1 yeast-extract malt extract (YEME) and tryptic soy broth (TSB) liquid media was used to cultivate mycelia for protoplast preparation and genome DNA isolation, and glucose-containing R5 agar media, with appropriate selective antibiotics, was used for protoplast regeneration after transformation. SFM (soy flour mannitol) agar was used for conjugation and cultivation of the spores. Phenotypic characterization was done on R5 medium and minimal media (MM) with 1% (w/v) mannitol supplemented with sugars as stated and, where appropriate, with the antibiotics apramycin (20 µg/ml) and/or thiostrepton (20 µg/ml) as selective markers.

All plasmids and oligonucleotides described in this work are summarised in Supplementary Table 1 and Supplementary Table 2 of the supplemental material, respectively. The shuttle vector pSET152[6] was used for genetic complementation and overexpression experiments, while the unstable multi-copy shuttle vector pWHM3[7] was exploited for gene replacement strategies[8]. Cre recombinase expressing plasmid, pUWLcre[9] was used for the creation of deletion mutants via genetic excision via *loxP* marked sites (see below for details). Expression vector pET-15b and pET-28a(+) (Novagen) were used for proteins heterologous expression. All DNA sequencing was performed by BaseClear BV (Leiden, The Netherlands).

#### Gene knock-out, complementation and overexpression

The detailed procedure for the creation of *S. coelicolor* gene replacement and deletion mutants is described previously[4]. Gene replacement mutants were generated via homologous recombination,

with the gene of interest replaced by the apramycin resistance cassette *aac(C)IV*. For this, the upstream and downstream flanking regions of *nagS* were PCR-amplified from *S. coelicolor* M145 genomic DNA using primer pairs SCO4393-LF/LR and SCO4393-RF/RR and cloned into pWHM3 using engineered *Eco*RI/*Xba*I and *Xba*I/*Hind*III restriction sites, respectively. The apramycin resistance cassette flanked by *loxP* sites was cloned in-between as an *Xba*I fragment. The resulting knock-out plasmid, designated pKO-4393, was introduced into *S. coelicolor* and Δ*nagB* via protoplast transformation. Correct recombination events were verified by appropriate antibiotics resistance and confirmed by PCR. To obtain deletion mutants, the apramycin resistance cassette was excised by introduction of the Cre recombinase expressing plasmid, pUWLcre[9], which allows for efficient removal of the cassette via the *loxP* recognition sites[10]. Deletion mutants were checked for the appropriate antibiotic sensitivity (loss of apramycin resistance) and confirmed by PCR.
For the complementation of *nagS*, DNA fragments of *nagS* coding region was amplified from the *S. coelicolor* genome DNA and cloned into pSET152, giving the vector pCOM-4393. NagS site-directed mutated sequences were amplified using two pairs of primers: 4393-compF/4393-compR and the corresponding mutated primers. The two amplified fragments were cloned into pSET152 by Gibson assembly[11], producing mutated *nagS* complementary vectors. The complementary constructs were introduced into *nagB-nagS* double mutant to obtain the corresponding complemented strain. For *nagS* and *nagA* overexpression, a 796-bp DNA fragment containing *nagS* coding region and a 1185-bp DNA fragment containing *nagA* coding region were amplified with the primers SCO4393-OE-F/R and SCO4284-OE-F/SCO4284-OE-R, and ligated simultaneously with the 310-bp highly efficient promoter *ermE*[12] into pSET152 to produce *nagS*-overexpressing vector pOE-4393 and *nagA*-overexpressing vector pOE-4284. The vectors were transformed into Δ*nagB* to generate *nagS* and *nagA* overexpressing strains, respectively.

#### Heterologous expression and purification of *S. coelicolor* NagA

For heterologous expression of *S. coelicolor* NagA in *E. coli*, the 1146-bp *nagA* coding regions was amplified from genome DNA using primer pair NagA-exp-F/NagA-exp-R. The PCR fragment was ligated into pET-28a (+) from *Xho*I and *Nco*I, generating expression vector pEX-4284. The NagA expression vector was transformed into *E. coli* BL21(DE3), and the expression of C-terminal $His_6$-tagged NagA recombinant protein were induced by addition of Isopropyl β-D-1-thiogalactopyranoside at the final concentration of 1.0 mM when the cell density reached around an optical density at 600 nm of 0.6. In addition, 1 mM $ZnCl_2$ was also supplemented into the cells as a range of NagA isozymes require divalent metal cations for their function[13,14], followed by incubation at 26°C overnight. Cells were harvested, washed, and disrupted in lysis buffer (50 mM sodium phosphate, 300 mM NaCl, 10 mM imidazole, pH 7.4) by sonication on ice. The preparations were then centrifuged, and the soluble $His_6$-tagged NagA from supernatant was purified using HisPur Cobalt Resin (Thermo fisher scientific; USA). After properly washing with wash buffer (50 mM sodium phosphate, 300 mM NaCl, 10 mM imidazole, pH 7.4), recombinant NagA was eluted from the resin with elution buffer (50 mM sodium phosphate, 300 mM NaCl, 300 mM imidazole, pH 7.4), and then desalted into phosphate buffer (25 mM phosphate pH 7.4, 100 mM NaCl, 1 mM DTT) and stored at -80°C.

#### Analytical size-exclusion chromatography

The oligomeric state of purified NagS was characterized by analytical size-exclusion chromatography. Protein at various concentrations (35 µM and 70 µM) was injected onto a Superdex 200 10/300 GL gel-filtration chromatography column (Cytiva) equilibrated in a buffer containing 20 mM HEPES pH 7.5, 300 mM NaCl, 5% glycerol, and 1 mM DTT. A calibration curve relating elution volume to molecular weight was generated by injecting a Protein Standard Mix (15 - 600 kDa; Sigma, product number: 69385) onto the same column with the same buffer.

#### Extinction co-efficient measurement of the product of NagS

To measure and calculate the extinction coefficient of NagS, an indirect method was used instead of spectrophotometry. As only a portion of the substrate is converted by NagS after the reaction, the concentration of the product is obtained by subtracting the final concentration of GlcNAc-6P, from the initial concentration of GlcNAc-6P (3.33 mM). First, the standard curve of the linear relationship between the concentrations of GlcNAc-6P (1 - 5 mM) and the peak areas of GlcNAc-6P measured by LC-MS was constructed (**Fig. S11**). The peak areas of GlcNAc-6P after the reaction were also measured (mean = 31467.63), thus the concentration of the remaining GlcNAc-6P was calculated as 2.15 mM using the standard equation. Thus, the concentration of NagS product was calculated as 1.18 mM (initial substrate concentration 3.33 mM minus final substrate concentration 2.15 mM). The final absorbance caused by the product was measured spectrophotometrically, which was 2.564. Therefore, the extinction co-efficient of the product of NagS was calculated as $2.17 \times 10^3$ $M^{-1} \cdot cm^{-1}$ through dividing the absorbance brought by the product by the concentration of it. All measurements involved were done in triplicate.

#### NagS-NagA coupled enzymatic assays

For NagS-NagA coupled assay, the reactions were carried out in the volatile buffer (50 mM *N*-ethylmorpholine/acetate pH 7.4), which is compatible with the subsequent Liquid

Chromatography/Mass Spectrometry (LC-MS) analysis. GlcNAc-6P (10 µL, 100 mM) was first incubated with NagS, followed by the addition of 150 nM NagA or heated-inactivated NagA after the NagS-catalysed reaction reached equilibrium. The reaction mixture was monitored at 230 nm until the absorbance stabilized, and then subjected to mass spectrometry analysis.

#### Thermal denaturation assays

The thermal stability and melting point of the protein was identified through measuring the intensity of SYPRO orange fluorescence. Upon the unfolding of the protein when temperature rises, the intensity of SYPRO orange increases due to the increased exposure of hydrophobic regions[15]. This change is monitored using a TOptical Real-Time qPCR Thermal Cycler (Biometra) or an iQ™5 Multicolor Real-Time PCR Detection System (BIO-RAD). NagS was tested for the thermal ability with or without 6-phosphogluconate (6-PG). For this, 200 µl of NagS at 1 µM was made up using the isolation HEPES buffer and 5X SYPRO orange (Invitrogen). A set of 3 repeats with a final volume of 50 µl of the protein solution per well was dispensed into a MicroAmp™ Optical 96-Well Reaction Plate (Applied Biosystems) and sealed by a Microseal® 'B' seal (BIO-RAD). The plate was then centrifuged at 2,000 rpm for 2 min at 10°C using an Allegra® X-15R centrifuge fitted with a µSX4250 rotor prior to the experiment. The level of SYPRO orange intensity was monitored between 20°C and 85°C. Additionally, 6-PG (1 mM and 5 mM) was tested to determine its possible effect on the thermal stability of NagS.

#### Inhibitor kinetic assays

The inhibitory activity of 6-phosphogluconate (6-PG) with respect to the enzymatic activity of NagS was determined. For this, the rate of reaction was measured by following the increase in UV absorbance at 230 nm as a function of time. Each assay (300 µL in total) was initiated by the addition of GlcNAc-6P (final concentration 0 - 8 mM) to a mixture containing 150 µL phosphate buffer (100 mM phosphate, 100 mM NaCl, pH 7.4), 6-PG (0.33 or 1 mM), and 220 nM NagS. Initial rates were analysed and fitted with the competitive inhibition model using GraphPad Prism (version 8.3.0). The rate equation used for $K_i$ calculation is $K_m' = K_m*(1+[I]/K_i)$.

#### Detection of compound 1-3 by LC-MS

For LC-MS analysis, the reactions were performed in a volatile buffer. In brief, 1 mM GlcNAc-6P was incubated in 50 mM *N*-ethylmorpholine acetate pH 7.4 at 30°C, and the reactions were initiated by the addition of 220 nM active NagS or boiled NagS (for the control sample). After the absorbance of the reaction mixtures had stabilised at 230 nm, the enzyme in the reaction mixture was precipitated with methanol and the supernatant obtained was then used for LC-MS analysis. LC-MS/MS acquisition was performed usinga Shimadzu Nexera X2 UHPLC system, with attached PDA, coupled to a Shimadzu 9030 QTOF mass spectrometer, equipped with a standard ESI source unit. A total of 2 µL of each sample was injected into a Waters Acquity HSS $C_{18}$ column (2.1 × 100 mm), which was run at a flow rate of 0.5 mL/min using 0.1% formic acid in $H_2O$ as solvent A, and 0.1% formic acid in acetonitrile as solvent B. The elution gradient used was 5% B for 1 min, 5–17% for 1 min, 17–20% for 8 min, then 20–100% for 1 min. All samples were analysed in negative polarity, using data dependent acquisition mode, in which full scan MS spectra (*m/z* 100–1700, scan rate 10 Hz, ID enabled) were followed by two data dependent MS/MS spectra (*m/z* 100–1700, scan rate 10 Hz, ID disabled) for the two most intense ions per scan at a collision energy of 20 eV. The parameters used for the ESI source were: interface voltage -3 kV, interface temperature 300°C, nebulizing gas flow 3 L/min, and drying gas flow 10 L/min.

### Suppemental Tables

#### Table S1 Bacterial strains and plasmids used in this study

| Bacterial strains | Description[*#] | Reference or source |
|---|---|---|
| *E. coli* | | |
| *E. coli* JM109 | General cloning strain | [1] |
| *E. coli* ET12567/pUZ8002 | Strain used for conjugation between *E. coli* and *Streptomyces* | [2] |
| *E. coli* Rosetta(DE3)pLysS | Strain used for protein expression | Novagen |
| *E. coli* BL21(DE3) | Strain used for protein expression | Novagen |
| *S. coelicolor* A3(2) | | |
| M145 | *S. coelicolor* A3(2) M145 SCP1- SCP2- prototroph | [3] |
| M145$^{E}$ | M145 complemented with empty pSET152 | This work |
| *ΔnagB* | M145 Δ*nagB*$^{d}$ | [4] |
| *ΔnagB*$^{E}$ | *ΔnagB* complemented with empty pSET152 | This work |
| *ΔnagAB* | M145 Δ*nagB*$^{d}$Δ*nagA*$^{d}$ | [4] |
| SMA11 | M145 Δ*nagB* suppressor, mutated in *nagS* | [5] |
| Δ*nagS* | M145 Δ*nagS*$^{d}$ | This work |
| Δ*nagB*Δ*nagS* | M145 Δ*nagB*$^{d}$Δ*nagS*$^{d}$ | This work |
| Δ*nagB*Δ*nagS*$^{C}$ | Δ*nagB*Δ*nagS* complemented with *nagS* | This work |
| Δ*nagB*Δ*nagS*$^{C\text{-}H53A}$ | Δ*nagB*Δ*nagS* complemented with *nagS* with mutation H53A | This work |
| Δ*nagB*Δ*nagS*$^{C\text{-}S54A}$ | Δ*nagB*Δ*nagS* complemented with *nagS* with mutation S54A | This work |
| Δ*nagB*Δ*nagS*$^{C\text{-}R64A}$ | Δ*nagB*Δ*nagS* complemented with *nagS* with mutation R64A | This work |
| Δ*nagB*Δ*nagS*$^{C\text{-}S91A}$ | Δ*nagB*Δ*nagS* complemented with *nagS* with mutation S91A | This work |
| Δ*nagB*Δ*nagS*$^{C\text{-}E94A}$ | Δ*nagB*Δ*nagS* complemented with *nagS* with mutation E94A | This work |
| Δ*nagB*Δ*nagS*$^{C\text{-}S119A}$ | Δ*nagB*Δ*nagS* complemented with *nagS* with mutation S119A | This work |
| Δ*nagB*Δ*nagS*$^{C\text{-}S121A}$ | Δ*nagB*Δ*nagS* complemented with *nagS* with mutation S121A | This work |
| Δ*nagB*Δ*nagS*$^{C\text{-}DD179A}$ | Δ*nagB*Δ*nagS* complemented with *nagS* with mutation D179A | This work |
| Δ*nagB*Δ*nagS*$^{C\text{-}N228A}$ | Δ*nagB*Δ*nagS* complemented with *nagS* with mutation N228A | This work |
| Δ*nagB*Δ*nagS*$^{E}$ | Δ*nagB*Δ*nagS* complemented with empty pSET152 | This work |
| Δ*nagS*$^{E}$ | Δ*nagS* complemented with empty pSET152 | This work |
| Δ*nagB-nagS*$^{OE}$ | Δ*nagB* with overexpressed *nagS* | This work |
| Δ*nagB-nagA*$^{OE}$ | Δ*nagB* with overexpressed *nagA* | This work |
| Δ*nagB*Δ*nagS-nagA*$^{OE}$ | Δ*nagB*Δ*nagS* with overexpressed *nagA* | This work |
| *ΔnagAB-nagS*$^{OE}$ | *ΔnagAB* with overexpressed *nagS* | This work |

| Plasmids | Description | Reference |
|---|---|---|
| pSET152 | Integrative *E. coli/Streptomyces* shuttle vector | [6] |
| pWHM3 | *E. coli/Streptomyces* shuttle vector, high copy number and unstable in *Streptomyces* | [7] |
| pET15b | Vector for His$_6$-tagged protein overexpression in *E. coli* | Novagen |
| pET-28a(+) | Vector for His$_6$-tagged protein overexpression in *E. coli* | Novagen |
| pUWLcre | *E. coli/Streptomyces* shuttle vector expressing the Cre recombinase in *Streptomyces* | [9] |
| pCOM-4393 | pSET152 harbouring *nagS* gene with its own promoter | This work |
| pCOM-4393-H53A | pSET152 harbouring *nagS* with the aa residue mutation H53A | This work |

| | | |
|---|---|---|
| pCOM-4393-S54A | pSET152 harbouring *nagS* with the aa residue mutation S54A | This work |
| pCOM-4393-R64A | pSET152 harbouring *nagS* with the aa residue mutation R64A | This work |
| pCOM-4393-S91A | pSET152 harbouring *nagS* with the aa residue mutation S91A | This work |
| pCOM-4393-E94A | pSET152 harbouring *nagS* with the aa residue mutation E94A | This work |
| pCOM-4393-S119A | pSET152 harbouring *nagS* with the aa residue mutation S119A | This work |
| pCOM-4393-S121A | pSET152 harbouring *nagS* with the aa residue mutation S121A | This work |
| pCOM-4393-D179A | pSET152 harbouring *nagS* with the aa residue mutation D179A | This work |
| pCOM-4393-N228A | pSET152 harbouring *nagS* with the aa residue mutation N228A | This work |
| pOE-4393 | pSET152 harbouring *nagS* gene under control of *ermE* promoter | This work |
| pOE-4284 | pSET152 harbouring *nagA* gene under control of *ermE* promoter | This work |
| pKO-4393 | pWHM3-oriT harbouring the *nagS* flanking regions with *loxP-aac(3)IV-loxP* | This work |
| pEX-4393 | pET15b harbouring the *nagS* gene | This work |
| pEX-4393-H53A | pET15b harbouring the *nagS* with the aa residue mutation H53A | This work |
| pEX-4393-S54A | pET15b harbouring the *nagS* with the aa residue mutation S54A | This work |
| pEX-4393-R64A | pET15b harbouring the *nagS* with the aa residue mutation R64A | This work |
| pEX-4393-S91A | pET15b harbouring the *nagS* with the aa residue mutation S91A | This work |
| pEX-4393-E94A | pET15b harbouring the *nagS* with the aa residue mutation E94A | This work |
| pEX-4393-S119A | pET15b harbouring the *nagS* with the aa residue mutation S119A | This work |
| pEX-4393-S121A | pET15b harbouring the *nagS* with the aa residue mutation S121A | This work |
| pEX-4393-D179A | pET15b harbouring the *nagS* with the aa residue mutation D179A | This work |
| pEX-4393-N228A | pET15b harbouring the *nagS* with the aa residue mutation N228A | This work |
| pEX-4284 | pET-28a(+) harbouring the gene *nagA* | This work |

* "d" indicates the gene before "d" is in-frame deleted; # "::*aac(3)IV*" indicates the gene before "::"is replaced by *aac(3)IV* cassette

#### **Table S2** Primers used in this study

| Name | 5'-3' sequence* | Function or descriptions |
|---|---|---|
| SCO4393-LF | GTCAGAATTCACGTCGATGCGCCGCGCCATAGG | *nagS* knock-out |
| SCO4393-LR | GAAGTTATCCATCACCTCTAGACTTGTGGTCGCTCATGCG | *nagS* knock-out |
| SCO4393-RF | GAAGTTATCGCGCATCTCTAGACGCCGCTGAACGCACCCGGTG | *nagS* knock-out |
| SCO4393-RR | GTCAAAGCTTGCGACGCTCCATTCGAGCAGAGG | *nagS* knock-out |
| SCO4393-KO-CF | CGTGCCCCTCGATGAGATTG | *nagS* mutant checking |
| SCO4393-KO-CR | ATGTTGCGCCGCTTGTAGAAC | *nagS* mutant checking |
| SCO4393-compF | CTATGACATGATTACGAATTCGATTGCCCTCGTCGGTCAGCTCC | *nagS* complementation |
| SCO4393-compR | TGGGCTGCAGGTCGACTACTGCTTCAGGCAGGTGAAGC | *nagS* complementation |
| SCO4393-H53A-F | TCGCCTTCGGCGCCGGCGCCTCCTCCCTCGCCGCCCAGG | NagS-H53A PCR |
| SCO4393-H53A-R | GCCGGCGCCGAAGGCGAAGAG | NagS-H53A PCR |
| SCO4393-R64A-F | GTCGTGTACGCCGCGGGCGGGCTCGCCCTG | NagS-R64A PCR |
| SCO4393-R64A-R | GCCCGCGGCGTACACGACGTCCTGGGCGGCGAG | NagS-R64A PCR |
| SCO4393-E94A-F | CTCGGCTCCGCCCTGGCGCGGGTCGACGGCCTCGCG | NagS-E94A PCR |
| SCO4393-E94A-R | CCAGGGCGGAGCCGAGGGTG | NagS-E94A PCR |
| SCO4393-S91A-F | ATGCCGGCCACCCTCGGCGCCGCCCTGGAGCGGGTCGACG | NagS-S91A PCR |
| SCO4393-S91A-R | GCCGAGGGTGGCCGGCATGAC | NagS-S91A PCR |
| SCO4393-S119A-F | GACGCCCTGGTGATCATCGCGCTCTCCGGGCGCAACG | NagS-S119A PCR |
| SCO4393-S119A-R | GATGATCACCAGGGCGTCGC | NagS-S119A PCR |
| SCO4393-D179A-F | TCCAAGATCGCCGTCGGCGCCGCGGAACTCACCCTCGACACC | NagS-D179A PCR |
| SCO4393-D179A-R | GCCGACGGCGATCTTGGAGTCC | NagS-D179A PCR |
| SCO4393-N228A-F | CCCGCTGCTGCGCTCGGGCGCCGTGGACGGCGGCCACGAATG | NagS-N228A PCR |
| SCO4393-N228A-R | CGAGCGCAGCAGCGGGGGTTC | NagS-N228A PCR |
| SCO4393-S54A-F | CTTCGGCGCCGGCCACGCCTCCCTCGCCGCCCAGGAC | NagS-S54A PCR |
| SCO4393-S54A-R | GTGGCCGGCGCCGAAGGCG | NagS-S54A PCR |
| SCO4393-S121A-F | CCTGGTGATCATCTCGCTCGCCGGGCGCAACGCCCTGCC | NagS-S121A PCR |

| | | |
|---|---|---|
| SCO4393-S121A-R | GAGCGAGATGATCACCAGGGCGTCG | NagS-S121A PCR |
| SCO4393-exp-F | GTACGAATTCATATGAGCGACCACAAGCCGGCC | NagS heterologous expression |
| SCO4393-exp-R | GTACGGATCCCCGGGGCACCGGGTGCGTTCA | NagS heterologous expression |
| NagA-exp-F | AATTTTGTTTAACTTTAAGAAGGAGATATACATGGCCCCAAGCAAGGTTCTCGC | NagA heterologous expression |
| NagA-exp-R | GTGGTGGTGGTGGTGCTCGCCCAGGTGGGGATCGACCAC | NagA heterologous expression |
| SCO4393-OE-F | ACTCCACAGGAGGACCCACAATGAGCGACCACAAGCCGGC | *nagS* overexpression |
| SCO4393-OE-R | GACCGAGCGTTCTGAACAAGTCAGCGGCGGTAGAAGATGCG | *nagS* overexpression |
| SCO4284-OE-F | CTCCACAGGAGGACCCACAATGGCCCCAAGCAAGGTTCTC | *nagA* overexpression |
| SCO4284-OE-R | GACCGAGCGTTCTGAACAAGTCAGCCCAGGTGGGGATCGAC | *nagA* overexpression |

* Underlined characters indicating restriction sites or overhangs for Gibson assembly. GAATTC, *Eco*RI; TCTAGA, *Xba*I; AAGCTT, *Hind*III.

#### Table S3 NagS data collection and model refinement statistics

| | Apo NagS | GlcNAc-6P bound NagS | 6-PG bound NagS |
|---|---|---|---|
| **Data collection** | | | |
| Space group | P 65 2 2 | P 65 2 2 | P 65 2 2 |
| Cell dimensions | | | |
| a, b, c (Å) | 87.69, 87.69, 273.49 | 88.37, 88.37, 284.46 | 87.67 87.67 278.81 |
| Α, β, γ (°) | 90, 90, 120 | 90, 90, 120 | 90, 90, 120 |
| Resolution (Å)* | 44.4 – 2.3 (2.38 – 2.30) | 45.7 - 2.6 (2.59 – 2.68) | 66.7 - 1.7 (1.72 - 1.69) |
| $R_{meas}$ | 0.167 (1.115) | 0.256 (2.918) | 0.068 |
| I / σI | 8.8 (1.7) | 9.7 (1.2) | 20.9 (1.7) |
| CC (1 / 2) | 0.993 (0.569) | 0.998 (0.527) | 1.000 (0.799) |
| Completeness (%) | 97.53 (95.97) | 99.90 (99.37) | 99.9 (98.6) |
| Multiplicity | 5.3 (5.1) | 18.9 (19.5) | 19.7 (20.4) |
| **Refinement** | | | |
| Resolution (Å) | 44.4 – 2.3 (2.38 – 2.3) | 45.7 - 2.6 (2.71 - 2.59) | 46.5 – 1.7 (1.71 – 1.69) |
| Number of reflections | | | |
| Used for refinement | 27893 (2646) | 21362 (2555) | 71889 (2724) |
| Used for $R_{free}$ | 1380 (124) | 1098 (128) | 3548 (144) |
| $R_{work}$ | 0.181 (0.254) | 0.191 (0.280) | 0.180 (0.347) |
| $R_{free}$ | 0.214 (0.306) | 0.229 (0.337) | 0.205 (0.354) |
| non-H atoms | 3887 | 3790 | 3853 |
| NagS | 3624 | 3626 | 3626 |
| ligands | 20 | 48 | 44 |
| water | 243 | 116 | 181 |
| R.m.s deviations | | | |
| Bond lengths (Å) | 0.006 | 0.002 | 0.011 |
| Bond angles (°) | 0.82 | 0.62 | 1.01 |
| Clash score | 2.2 | 1.9 | 1.4 |
| Ramachandran | | | |
| Favored | 98.0% | 99.0% | 99.0% |
| Outliers | 0.3% | 0.4% | 0.4% |
| Average B-factor | 35 | 61 | 45 |
| Protein | 35 | 61 | 45 |
| Ligands | 50 | 69 | 44 |
| Water | 37 | 54 | 44 |
| PDB ID code | 9F7O | 9F7V | 9EOL |

* Values in parentheses are for highest-resolution shell.

#### Table S4 NMR data of 1 as compared to Chromogen I[16]

| Position | | $\delta_H$, mult. (*J* in Hz) | | $\delta_C$, type | |
|---|---|---|---|---|---|
| | | **1**[a] | Chromogen I[b] | **1**[c] | Chromogen I[d] |
| α-anomer[e] | 1 | 6.06, dd (4.0, 1.0) | 6.01, dd (4.0, 0.9) | 102.1, CH | 102.4, CH |
| | 2 | | | ND | 137.3, C |
| | 3 | 6.19, dd (1.7, 1.0) | 6.14, br s | 110.4, CH | 112.2, CH |
| | 4 | 5.10, td (4.0, 1.7) | 5.03, td (4.0, 1.6) | 87.7, CH | 87.9, CH |
| | 5 | 3.89, m | 3.80, td (4.0, 7.2) | 75.6, CH | 76.4, CH |
| | 6 | a: 3.88, m<br>b: 3.79, m | a: 3.69, dd (11.9, 4.0)<br>b: 3.55, dd (11.9, 7.2) | 67.8, $CH_2$ | 65.4, $CH_2$ |
| | $CH_3$ (*N*-acetyl) | 2.11, s | 2.10, s | 25.4, $CH_3$ | 25.6, $CH_3$ |
| | CO (*N*-acetyl) | | | 176.2, C | 176.5, C |
| β-anomer | 1 | 5.99, t (1.0) | 5.97, d (1.0) | 102.0, CH | 102.3, CH |
| | 2 | | | ND | 136.8, C |
| | 3 | 6.22, dd (1.4, 1.0) | 6.20, dd (1.6, 1.0) | 113.3, CH | 113.1, CH |
| | 4 | 4.88, m | 4.81, d (1.6) | 87.2, CH | 87.5, CH |
| | 5 | 3.81 | 3.75–3.71, m | 76.0, CH | 76.8, CH |
| | 6 | 3.93, m | a: 3.75–3.71, m<br>b: 3.64–3.59, m | 67.9, $CH_2$ | 65.5, $CH_2$ |
| | $CH_3$ (*N*-acetyl) | 2.11, s | 2.10, s | 25.4, $CH_3$ | 25.6, $CH_3$ |
| | CO (*N*-acetyl) | | | 176.2, C | 176.5, C |

[a] 600 MHz at 298 K in $D_2O$, [b] 500 MHz in $D_2O$. Temperature is not given, [c] 213 MHz at 298 K in $D_2O$, [d] 125 MHz in $D_2O$. Temperature is not given, [e] Ratio of the anomers α:β is 1:0.6 for both **1** and chromogen I, ND: Not detectable

#### Table S5 HRMS data of the compounds identified in this study

| Compounds | Molecular formula | Calculated $[M-H]^-$ *m/z* | Observed *m/z* |
|---|---|---|---|
| **1** | $C_8H_{14}NO_8P$ | 282.0384 | 282.0387 |
| **2/3** | $C_6H_{12}NO_7P$ | 240.0279 | 240.0280 |

#### Supplemental Figures

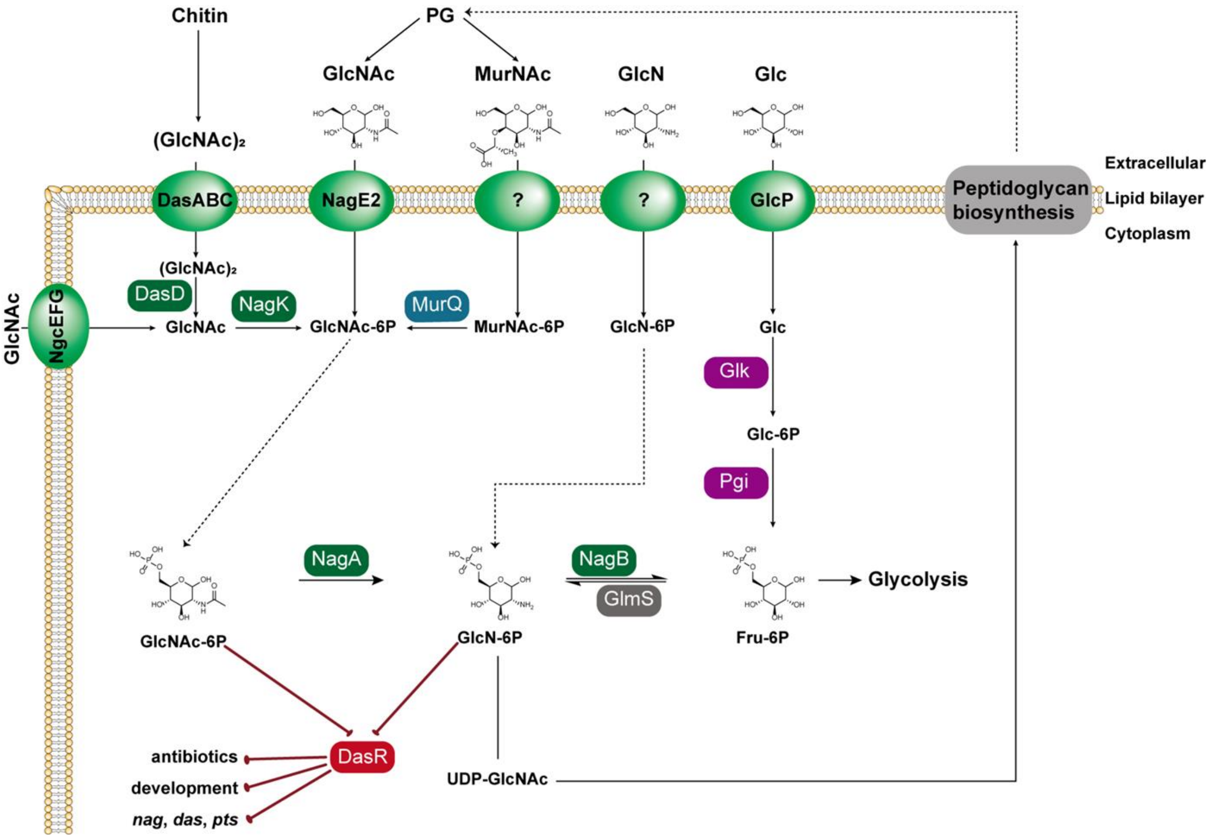

**Figure S1. Metabolic pathway of aminosugar in *Streptomyces*.** Peptidoglycan degradation releases monomers of GlcNAc and MurNAc, which are subsequently taken up by the cells for recycling. The phosphoenolpyruvate-dependent phosphotransferase system (PTS) phosphorylates monomeric GlcNAc during transport into GlcNAc-6P. Subsequently, GlcNAc-6P is metabolized by NagA and NagB to Fru-6P, which enters into glycolysis. Limited information is available on GlcN transport and metabolism in *S.coelicolor*. Besides its metabolism to fructose-6P, GlcN-6P is also the starting point for the biosynthesis of Lipid II, the building block for cell-wall synthesis. GlcNAc-6P and GlcN-6P are effector molecules for DasR, which is a global repressor of among others aminosugar metabolism, natural product biosynthesis and development in *Streptomyces*. Metabolic routes are represented by arrows with corresponding enzymes. For clarity, the substrates and enzyme names are abbreviated. Abbreviations not mentioned in the text: Glk, glucokinase; Pgi, glucose-6-phosphate isomerase; DasD, *N*-acetyl-β-D-glucosaminidase.

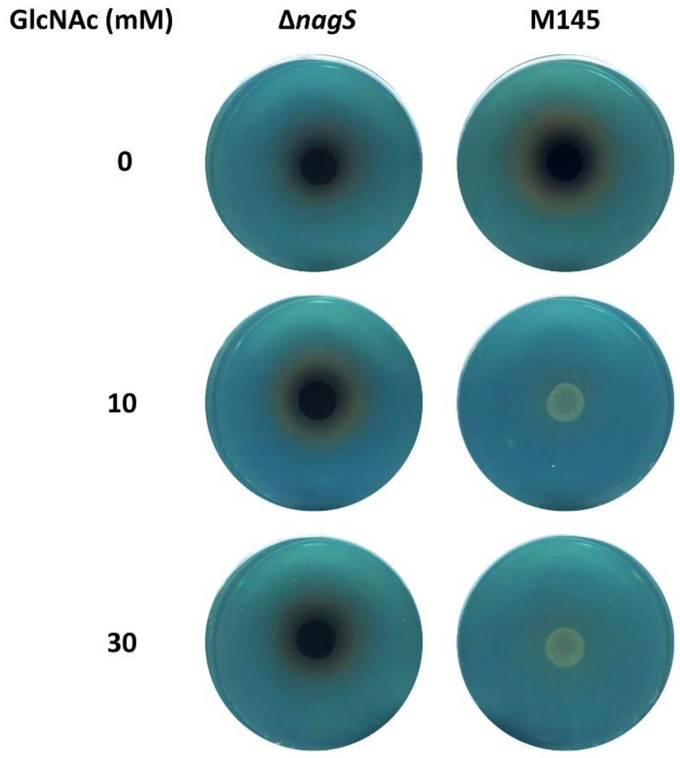

**Figure S2**. **Inhibition of siderophore production by GlcNAc requires NagS.**

Streptomycetes were grown as spots on R5 agar plates by plating 10 µl of $10^8$ spores/ml and after incubation at 30°C overnight, plates were overlaid with Chrome azurol S (CAS) staining solution and examined visually. Larger orange halos show siderophore production, dark central circles are caused by the pigmented anibiotic actinorhodin. Note that the biosynthesis of siderophores and antibiotic is not repressed by GlcNAc in *nagS* mutants. CAS assays were carried out as described previously[17].

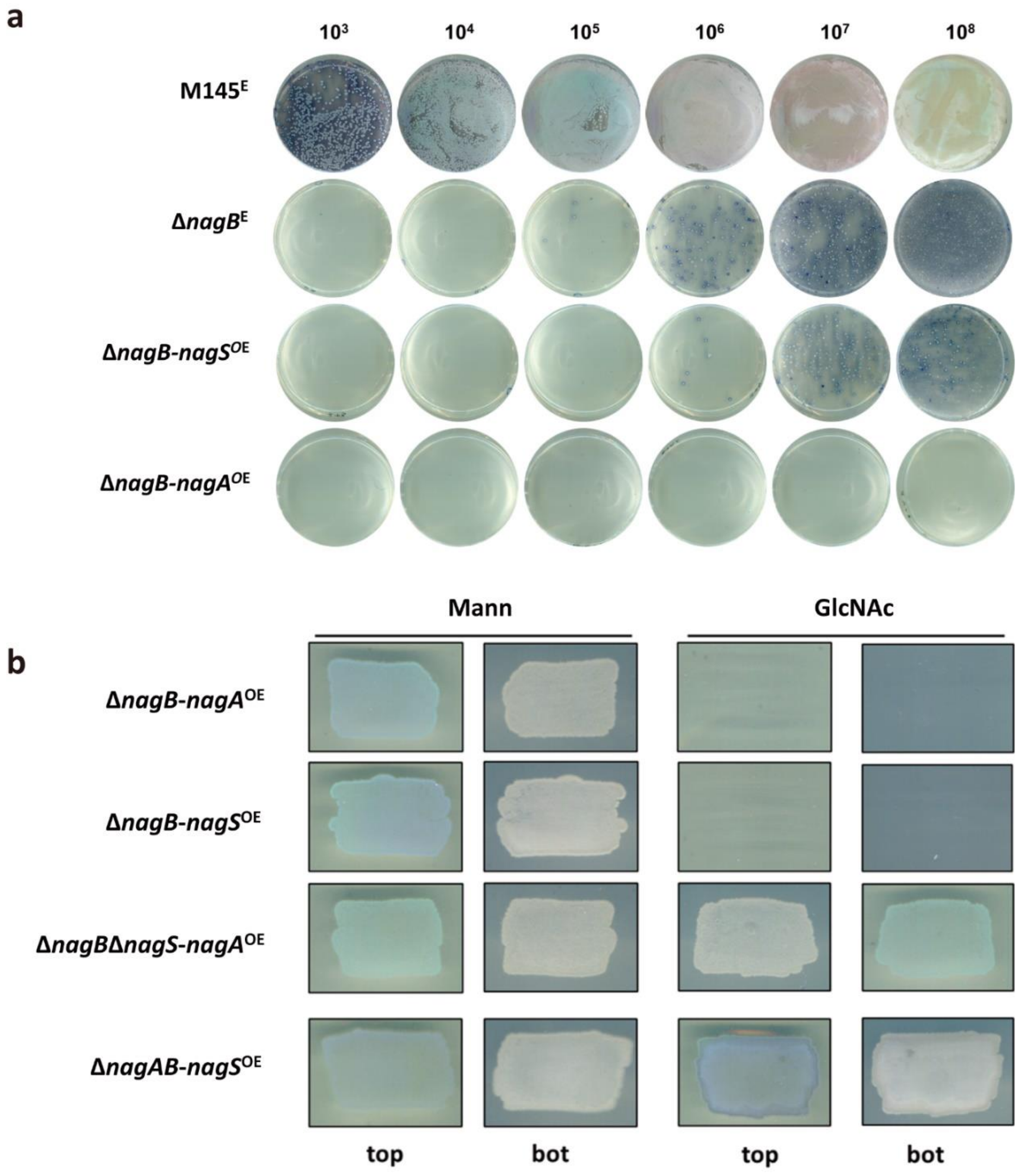

**Figure S3. Roles of NagS and NagA in GlcNAc toxicity.** (**a**) Suppressor mutants check for *nagB* with overexpressed *nagS* and *nagA*. Spores with different CFU of M145 complemented with empty pSET152 (M145$^{E}$), Δ*nagB* complemented with empty pSET152 (Δ*nagB*$^{E}$), Δ*nagB* complemented with *nagS* expressed by *ermE* (Δ*nagB-nagS*$^{OE}$), and Δ*nagB* complemented with *nagA* expressed by *ermE* (Δ*nagB-nagA*$^{OE}$) were streaked on the MM supplemented with 1% mannitol and 10 mM GlcNAc. After 72 h-culturing, the numbers of suppressor mutants were compared. (**b**) Effect of overexpression of *nagA* and *nagS* on the sensitivity of *S. coelicolor nagB* mutants to GlcNAc. Spores ($5 \times 10^5$ CFU) of Δ*nagB* with overexpressed *nagA* (Δ*nagB-nagA*$^{OE}$), Δ*nagB* with overexpressed *nagS* (Δ*nagB-nagS*$^{OE}$), Δ*nagB*Δ*nagS* with overexpressed *nagA* (Δ*nagB*Δ*nagS-nagA*$^{OE}$), and Δ*nagAB* with overexpressed *nagS* (Δ*nagAB-nagS*$^{OE}$) were streaked onto MM with 1% mannitol (Mann), or with 1% mannitol and 10 mM GlcNAc (GlcNAc). The strains were cultured for 72 h at 30°C.

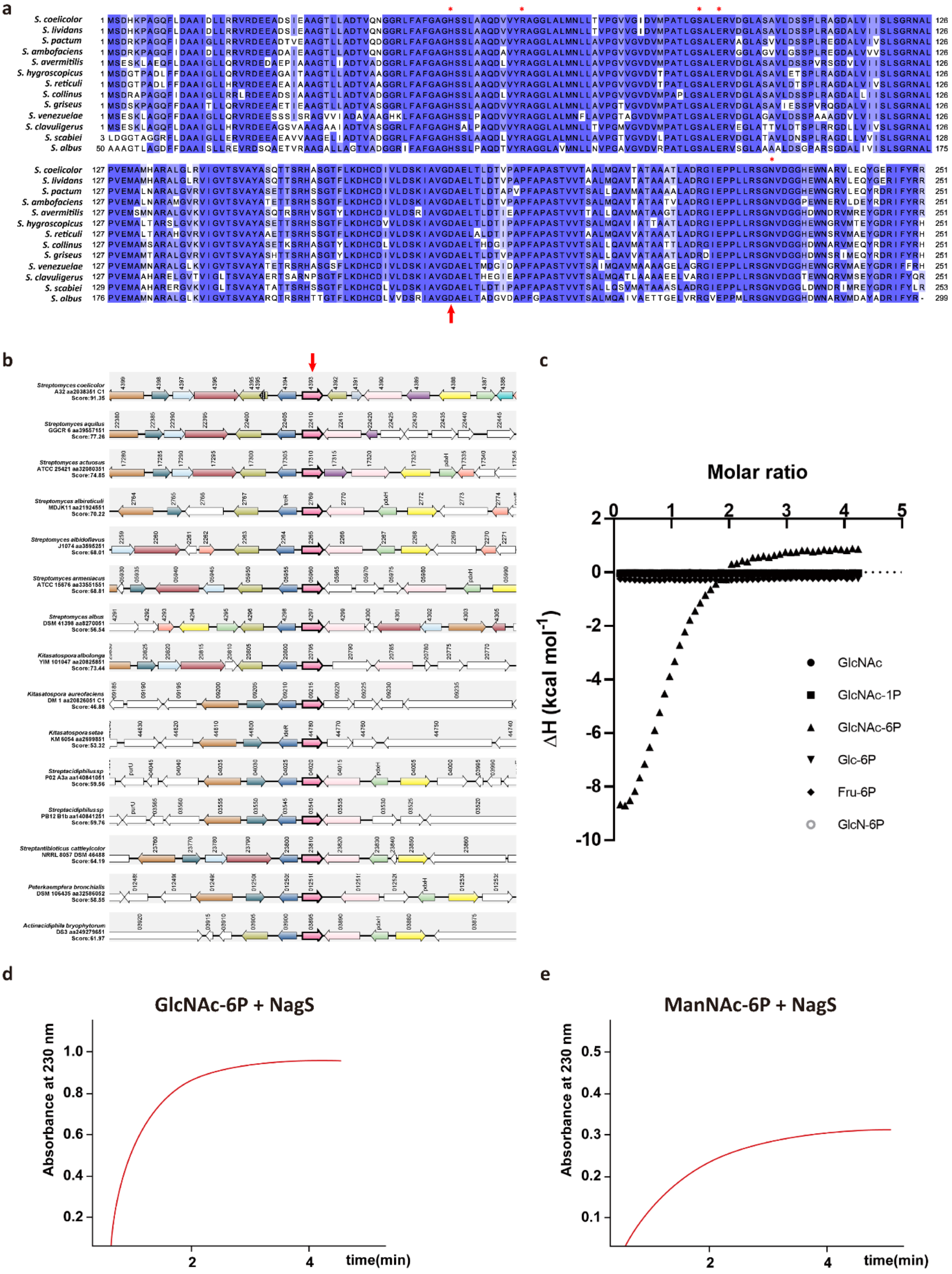

**Figure S4. NagS is a conserved and novel enzyme involved in aminosugar metabolism.** (a) Alignment of NagS protein sequence with its homologs from other *Streptomyces* species up to residue 240. Identical amino acids are shown in dark blue, and amino acids with similar properties in light blue. The D179N mutation identified in SMA11 is indicated with the red arrow below. Residues are determined to be important for catalysis are indicated with red stars above. Alignments was analysed by Clustal Omega and the image was generated using Jalview (Version 2.11.2.7). (**b**) Gene synteny of *nagS* (SCO4393) and its homologs in other Streptomycetaceae. Note that *nagS-dmdR1* is conserved in all *Streptomycetaceae* family except *Yinghuangia* genus. Analysis was done by Synttax inputting NagS aa sequence and the scores are given. Homologous genes are presented in the same colours with *nagS* homologous genes indicated by the red arrow. (**c**) Initial ITC study of NagS. For ITC binding studies, 1 mM ligand was titrated with 6 or 8 µL injections into 50 mM purified NagS. Fru-6P, Glc-6P, GlcN-6P, GlcNAc, GlcNAc-1P, and GlcNAc-6P were tested. Note that NagS was only able to bind GlcNAc-6P. Absorbance changes detected at 230 nm when incubating 2 mM GlcNAc-6P (**d**) or ManNAc-6P (**e**) with NagS at 30°C.

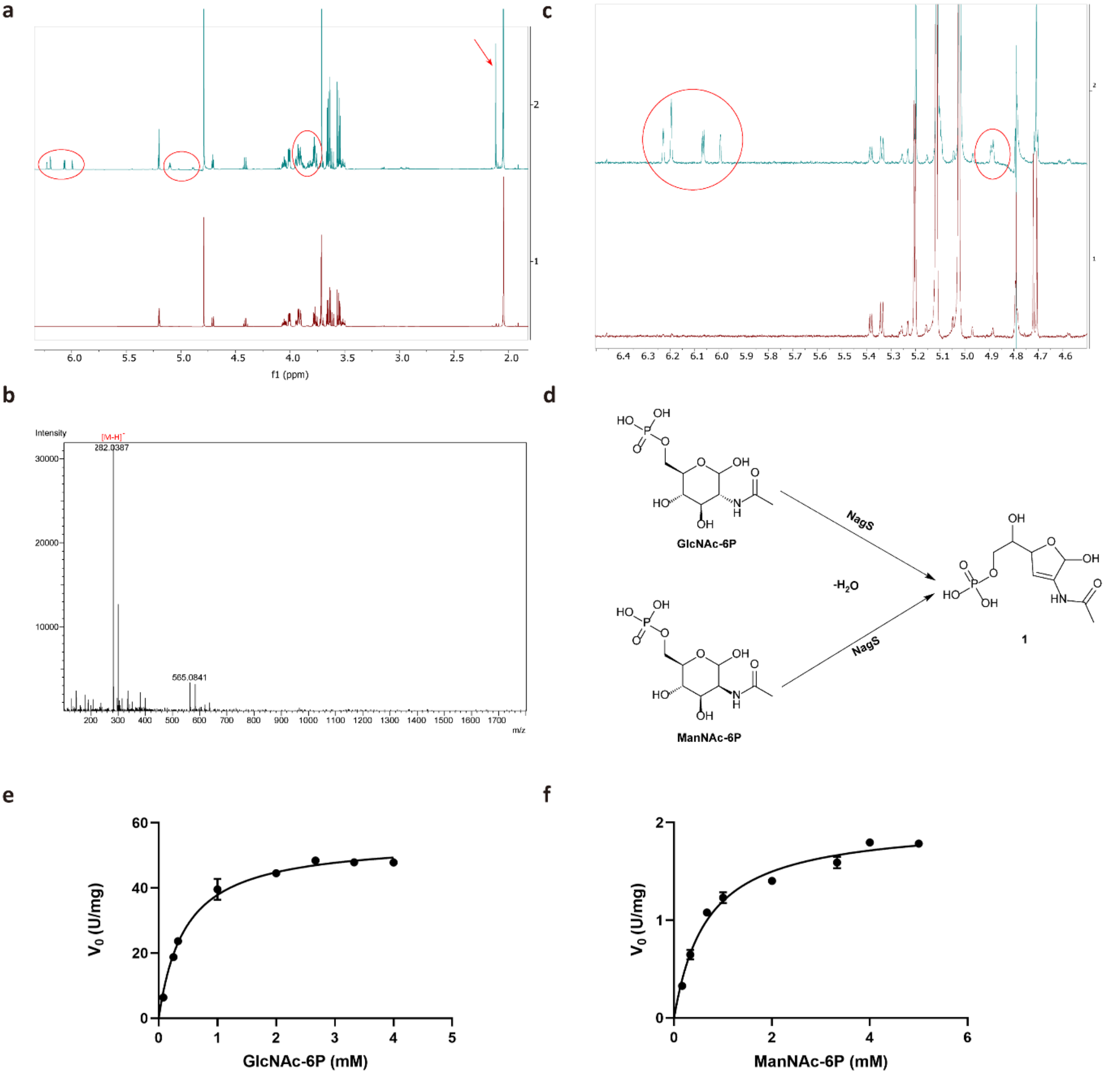

**Figure S5. Function determination and kinetics of NagS.** (**a**) $^1$H NMR spectrum of the enzymatic reaction mixture of GlcNAc-6P with either the active NagS (top) or the heated-inactivated one (bottom). The associated NMR peaks of the reaction product are highlighted in red circles and arrows. (**b**) HRESIMS spectrum of the NagS product, compound **1**: *m/z* 282.0387 [M-H]$^-$ (calculated for $C_8H_{13}NO_8P$, 282.0384) (**c**) $^1$H NMR spectrum of the enzymatic reaction mixture of ManNAc-6P with either the active NagS (top) or the heat-inactivated one (bottom). The associated NMR peaks of the reaction product are highlighted in red circles. (**d**) Reactions catalysed by NagS. NagS dehydrates both GlcNAc-6P and ManNAc-6P to produce compound **1**. Michaelis-Menten curves were fitted, and selected curves are shown for NagS with the substrates GlcNAc-6P (**e**) with $K_m$ value of 0.45 ± 0.03 mM and $k_{cat}/K_m$ value of 5.48 × 10$^4$ M$^{-1}$·s$^{-1}$, and ManNAc-6P (**f**) with $K_m$ value of 0.68 ± 0.05 mM and $k_{cat}/K_m$ value of 1.32 × 10$^3$ M$^{-1}$·s$^{-1}$. In **e** and **f**, the $V_0$ data were plotted against the substrate concentration, and each assay was performed in triplicate and expressed as a mean ± standard error.

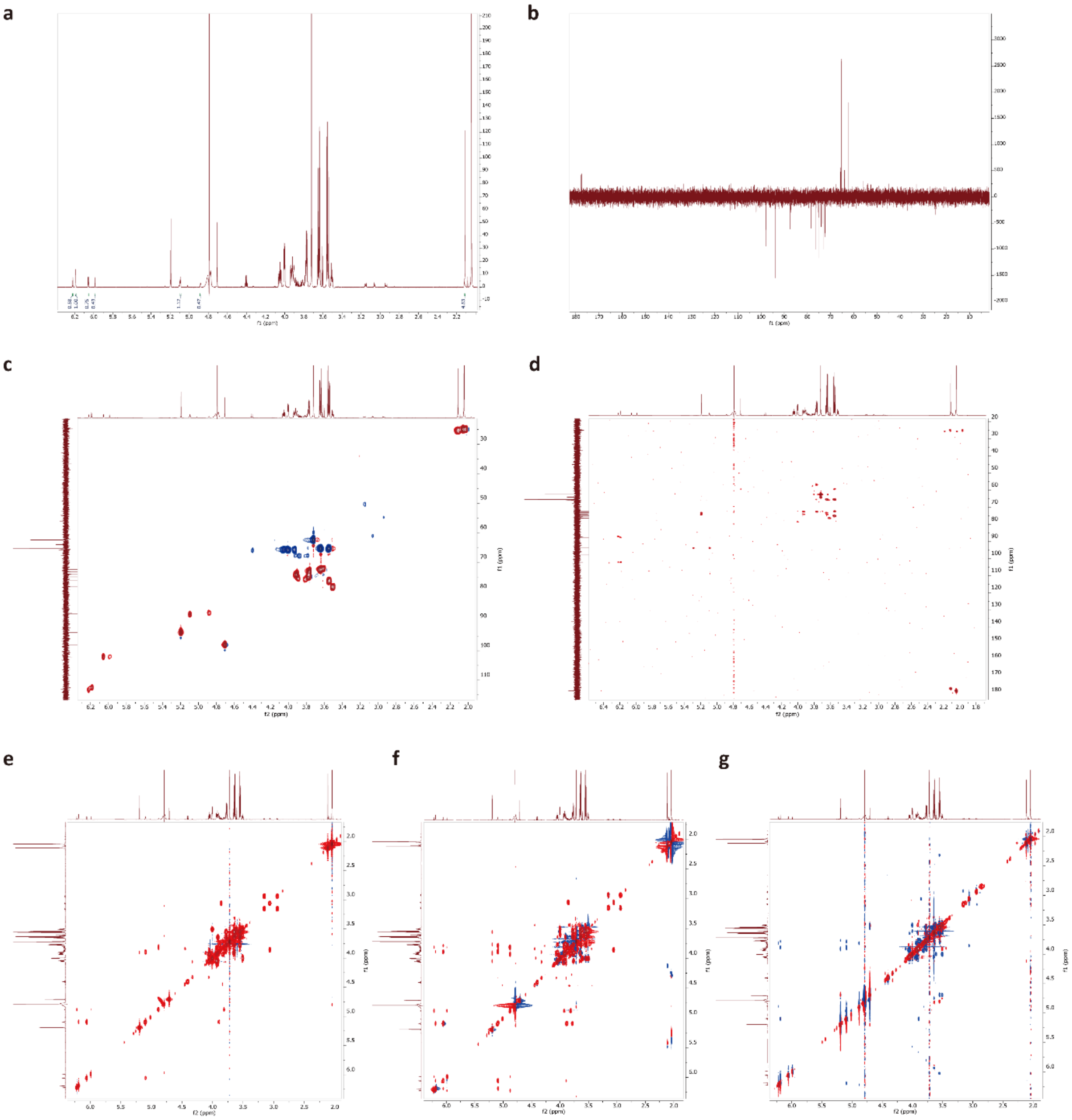

**Figure S6. Spectroscopic data of compound 1.** (**a**) $^{1}$H NMR spectrum of **1** in the reaction mixture (600 MHz, in $D_2O$). Non-overlapping peaks are integrated. (**b**) $^{13}$C NMR spectrum of **1** in the reaction mixture (213 MHz, in $D_2O$). (**c**) Multiplicity-edited HSQC spectrum of **1** in the reaction mixture (600 MHz, in $D_2O$). (**d**) HMBC spectrum of **1** in the reaction mixture (600 MHz, in $D_2O$). (**e**) COSY spectrum of **1** in the reaction mixture (600 MHz, in $D_2O$). (**f**) TOCSY spectrum of **1** in the reaction mixture (600 MHz, in $D_2O$). (**g**) NOESY spectrum of **1** in the reaction mixture (600 MHz, in $D_2O$).

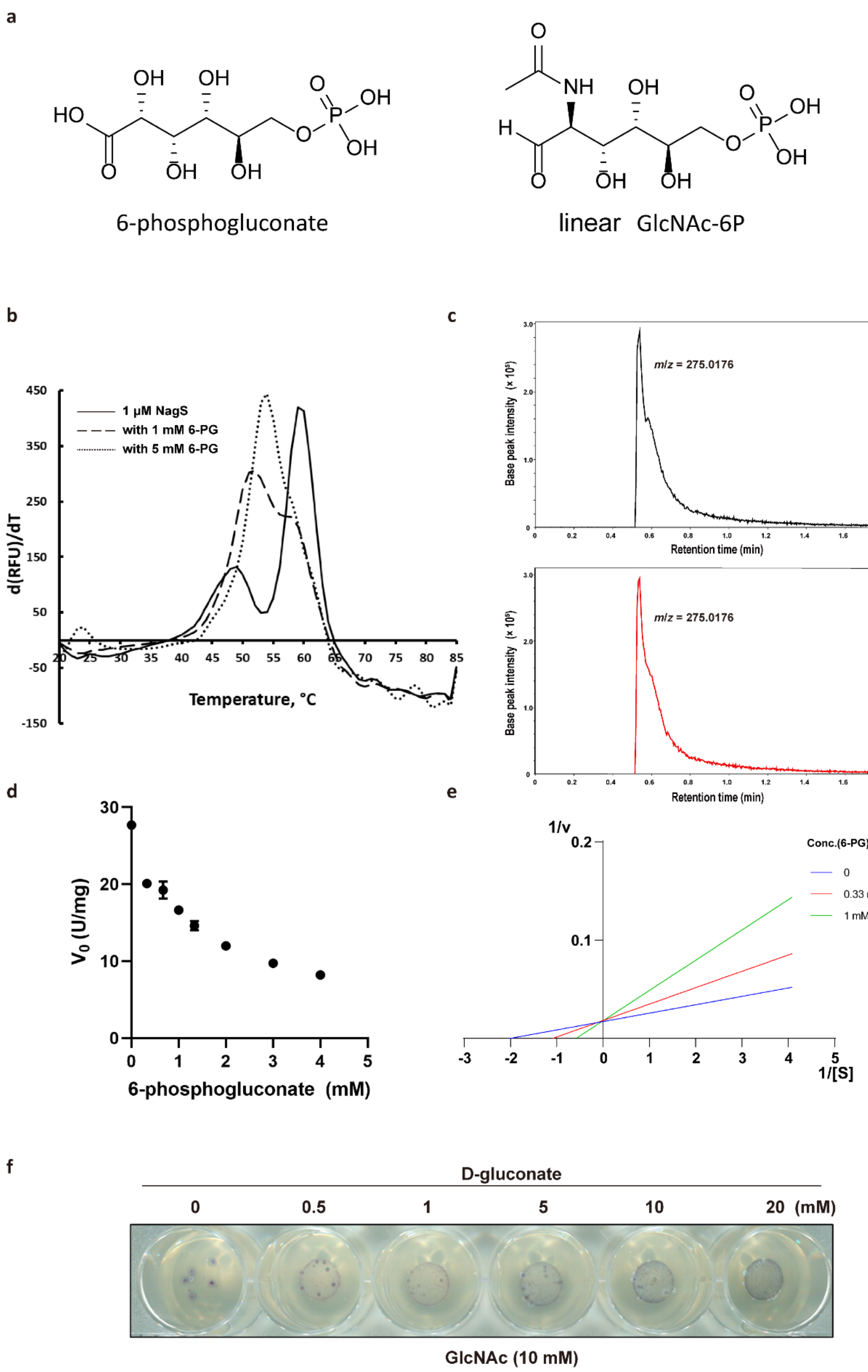

**Figure S7. Inhibitory effect of 6-phosphogluconate (6-PG).** (**a**) Chemical structures of 6-PG and linear GlcNAc-6P. (**b**) Average $T_m$ curve of NagS in presence of 6-PG. Melting curve of 1 µM NagS (black), 1 µM NagS with 1 mM 6-PG (dash), and 5 mM 6-PG (dot). It was shown that when 6-PG concentration increases, a shift in $T_m$ was shown. The two $T_m$ peaks at 48.84°C ± 0.10°C and 59.53°C ± 0.31°C merged at into a single peak at $T_m$ 53.95°C ± 0.20°C in response to the addition of 5 mM 6-

PG. (**c**) 6-PG detection by LC-MS. Reactions of 10 mM 6-PG with NagS (black) or deactivated NagS (red) were detected by LC-MS. Note that no 6-PG (*m*/*z* = 275.0176) was consumed in both conditions. (**d**) Evaluation of the inhibition of 6-PG on NagS activity. The activity of NagS ($V_0$) was measured using 1 mM GlcNAc-6P as the substrate, with the addition of 0-4 mM 6-PG. (**e**) Competitive inhibition of NagS by 6-PG. The inhibition by 6-PG is presented as Lineweaver-Burk plot ($K_i$ = 0.28 mM). (**f**) Effect of the addition of D-gluconate on GlcNAc sensitivity. Spores ($5 \times 10^5$ CFU) suspension of *S. coelicolor* M145 *nagB* mutant was spotted on MM supplemented with 1% mannitol, 10 mM GlcNAc and a range concentration of D-gluconate, followed by incubation for 72 h at 30°C.

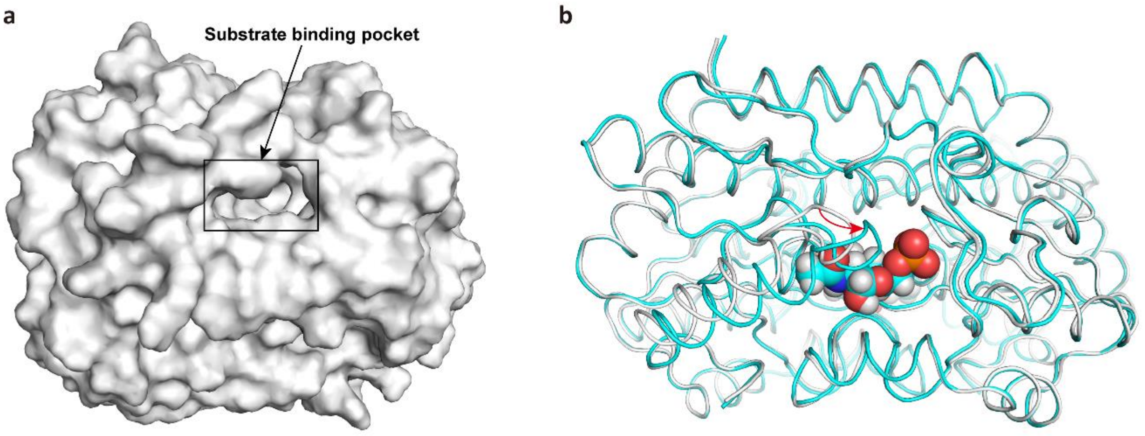

**Figure S8. Substrate binding pocket and the movable loop of NagS**. (**a**) Substrate binding pocket of NagS located at the dimeric interface. (**b**) Secondary structure alignment of ligand-free (white) and ligand-bound (cyan) NagS. The substrate GlcNAc-6P is shown in the form of sphere. The direction and angle of movement of this loop after binding the substrate is indicated by the red arrow. Note that this loop moves towards the bound substrate after binding to it.

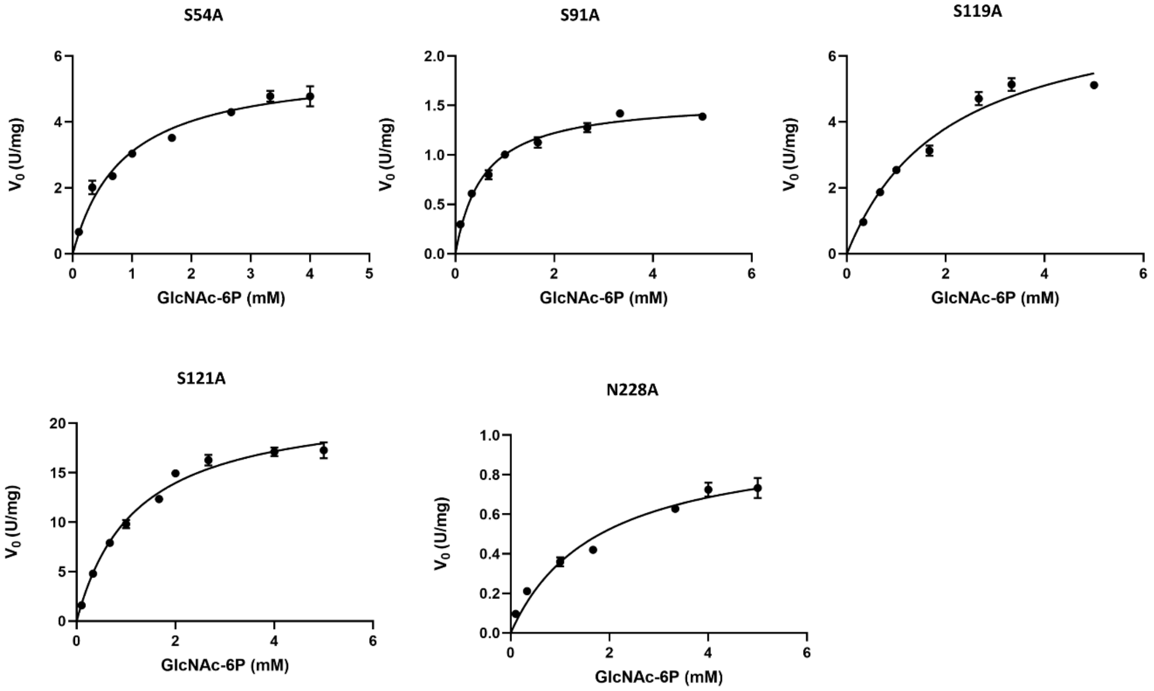

**Figure S9. Kinetics of NagS variants**. Michaelis-Menten curves were fitted, and selected curves are shown for NagS variants with the substrates GlcNAc-6P. The $V_0$ data were plotted against the substrate concentration, and each assay was performed in triplicate and expressed as a mean ± standard error.

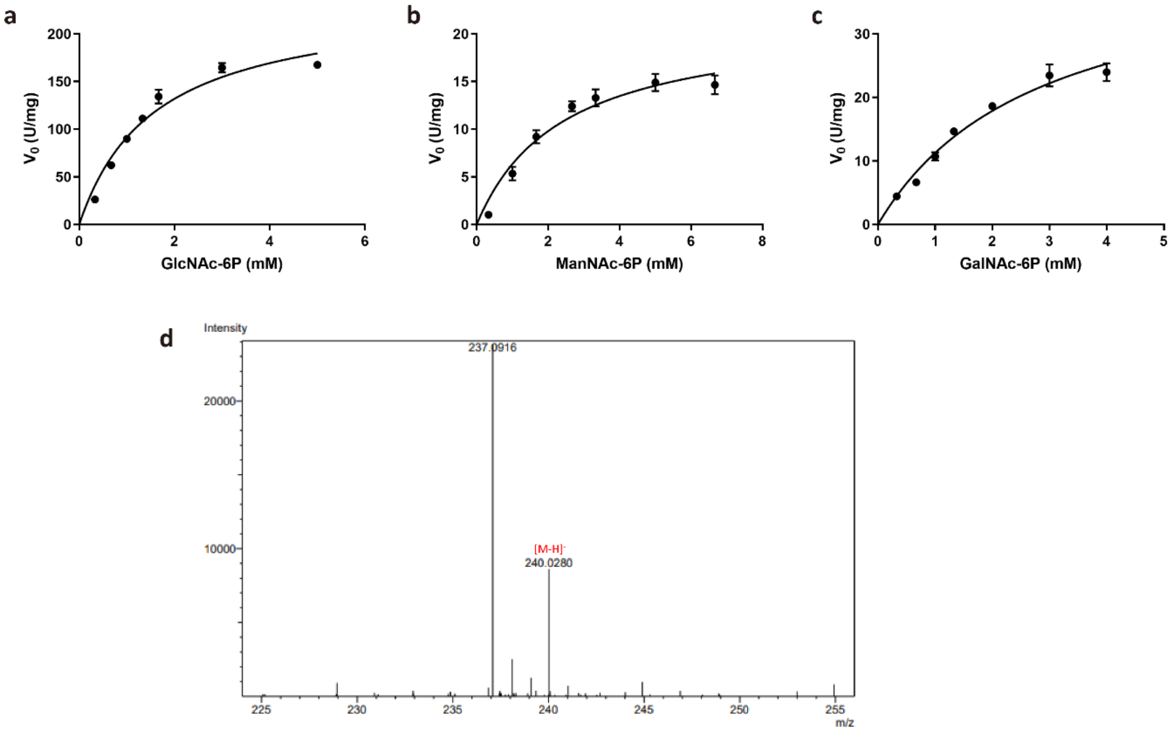

**Figure S10. Kinetics of *S. coelicolor* NagA and the product identification of NagS-NagA catalysis.** Michaelis-Menten curves were fitted, and selected curves are shown for *S. coelicolor* NagA with the substrates: GlcNAc-6P (**a**) with $K_m$ value of 1.59 ± 0.21 mM and $k_{cat}/K_m$ value of 1.01 × $10^5$ $M^{-1}·s^{-1}$, ManNAc-6P (**b**) with $K_m$ value of 2.40 ± 0.44 mM and $k_{cat}/K_m$ value of 6.06 × $10^3$ $M^{-1}·s^{-1}$, and GalNAc-6P (**c**) with $K_m$ value of 2.84 ± 0.41 mM and $k_{cat}/K_m$ value of 1.03 × $10^4$ $M^{-1}·s^{-1}$. (**d**) HRESIMS spectrum of the compound **2**/**3**: *m*/*z* 240.0280 $[M-H]^-$ (calculated for $C_6H_{12}NO_7P$, 240.0279). In **a**-**c**, the $V_0$ data were plotted against the substrate concentration, and each assay was performed in triplicate and expressed as a mean ± standard error.

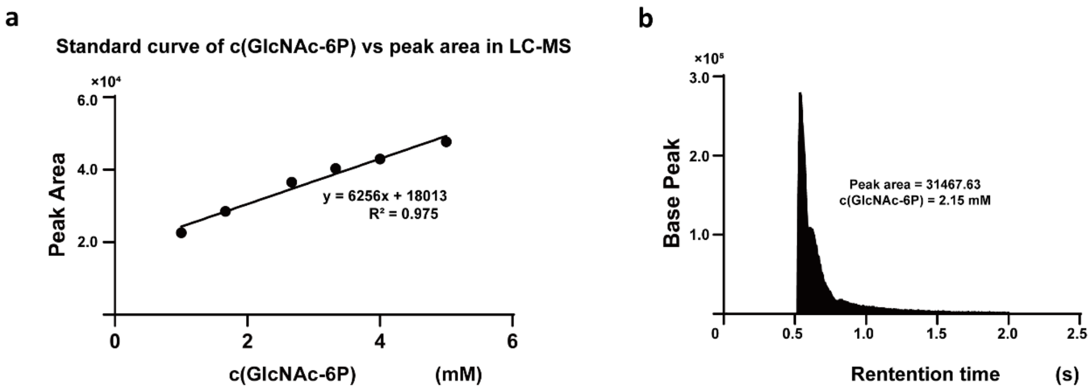

**Figure S11. Extinction co-efficient measurement of NagS.** (**a**) This standard curve of GlcNAc-6P was obtained by plotting the concentration of GlcNAc-6P (1, 1.67, 2.67, 3.33, 4 and 5 mM) with the corresponding peak area detected in the LC-MS spectrum. (**b**) Area of the GlcNAc-6P peak after the reaction catalysed by NagS. The concentration of remaining GlcNAc-6P is calculated to be 2.15 mM.
